# Pseudospatial transcriptional gradient analysis of hypothalamic ependymal cells: towards a new tanycyte classification

**DOI:** 10.1101/2023.07.06.547914

**Authors:** M. Brunner, D. Lopez-Rodriguez, A. Messina, B. Thorens, F Santoni, F. Langlet

**Author notes:** co-corresponding authors, Address: Department of Biomedical Sciences, University of Lausanne, Bugnon 7, 1005 Lausanne, Switzerland., Contact 1, Contact 2. Co-First (M. Brunner, D. Lopez-Rodriguez).

## Abstract

The ependyma lining the third ventricle (3V) in the mediobasal hypothalamus is recognized as a critical player in controlling energy balance and glucose homeostasis. Its molecularly distinct cell types, including diverse tanycyte subpopulations and typical ependymal cells, confer a high functional heterogeneity. The study of gene expression profiles and dynamics of ependymal cells has the potential to uncover fundamental mechanisms and pathways involved in metabolic regulation.

Here, we cataloged 5481 hypothalamic ependymocytes using FACS-assisted single-cell RNA sequencing from fed, 12h-fasted, and 24h-fasted adult male mice. First, standard clustering analysis revealed the limitation of the current characterization regarding the different ependymal cell subpopulations along the 3V. Indeed, while typical ependymal cells and β2-tanycytes are sharply defined at the molecular level, other subpopulations (*i.e*., β1-, α2-, and α1 tanycytes) display fuzzy boundaries and very few specific markers. Moreover, we observed that 12h- and 24h-fasting dynamically modulate gene expression, increasing tanycyte subgroup heterogeneity. Secondly, pseudospatial trajectory analysis based on peculiar UMAP neuroanatomical distribution improved the identification of tanycyte markers, distinguishing specific *versus* overlapping features and better segregating tanycyte specific *versus* standard functions. Intriguingly, we discovered numerous functions related to tanycyte-neuron and tanycyte-synapse interactions with modulation by energy balance. Finally, combining pseudospatial analysis and gene regulatory network inference, we observed that fasting dynamically shifts patterns in gene expression and transcription activity along the 3V, creating a metabolic and functional switch for some subpopulations.

Altogether, this data provides a mechanism through which energy status leads to distinct cell type-specific responses along the 3V and gives new insights into molecular diversity underlying tanycyte classification.

## INTRODUCTION

The central nervous system comprises numerous neural cells with a high morphological, molecular, and functional diversity, interconnected in precise and dynamic networks to ensure countless physiological functions. Among these functions, energy balance is mainly orchestrated by hypothalamic and brainstem circuits^1, 2^. Neurons and glial cells, including astrocytes, tanycytes, and microglia, compose these circuits and regulate food intake, energy expenditure, and glucose homeostasis^3–5^.

Among glial cells, ependymocytes lining the third ventricle (3V) are now recognized full-fledged actors in regulating energy balance and glucose homeostasis^6, 7^. The striking feature of the ependyma in the mediobasal hypothalamus (MBH) is its high heterogeneity, conferring various biological functions^8^. First, lining the top of the 3V, typical ependymal cells are cuboid and ciliated-epithelial glial cells that control cerebrospinal fluid (CSF) homeostasis and waste clearance^9^. A coordinated ciliary beating of these cuboidal cells allows the maintenance of CSF flows through the ventricular system^10^. Interestingly, the metabolic peptide melanin-concentrating hormone increases cilia beat frequency, increasing CSF flow and volume transmission to possibly meet metabolic needs^11^. Besides typical ependymal cells, tanycytes are polarized ependymocytes lining the bottom and the lateral walls of the 3V in the tuberal region of the hypothalamus^7, 12^. Their unique morphology and location allow them to form a triple interface between the ventricular system, the brain parenchyma, and the vascular compartment for regulating energy balance^13–15^. Indeed, their cell bodies lining the ventricular wall favor sensing the central metabolic state in the CSF^23, 25^. Their long cytoplasmic extensions into key hypothalamic brain areas -including the median eminence (ME), the arcuate nucleus (ARH), the ventromedial nucleus (VMH), and the dorsomedial nucleus (DMH)-enable the integration into circuits regulating energy balance^15^. Finally, their endfeet contacting vessels in these brain areas allow the reception of metabolic information from the bloodstream^16, 20, 37^. Besides this strategic location, tanycytes are highly heterogeneous, with distinct gene expression, morphological, and functional properties^7, 8^. In the context of energy balance regulation, tanycytes have been described as regulators of blood-brain^16, 17^ and blood-CSF^18–20^ exchanges, transporters of peripheral metabolic signals into the brain^19, 20^, controllers of neurosecretion into the peripheral circulation^21^, detectors of the individual’s metabolic state^22–24^, coordinators of neuronal functions^25–27^, and modulators of neural circuits through their neural stem cell properties^28–30^.

Thus, based on their location along the 3V and these functional disparities, tanycytes are historically classified into four subtypes: β1, β2, α1, and α2^31, 32^. β2 tanycytes line the floor of the 3V in the median eminence, contact the fenestrated vessels of the hypothalamic-hypophysial portal system, and mainly regulate blood-brain exchanges. β1 tanycytes line the lateral evaginations of the infundibular recess and the ventromedial arcuate nucleus (vmARH), contact *en-passant* different neural cells, including neurons, before ending on the lateral ME fenestrated vessels or the pial surface of the brain. α2 tanycytes line the dorsomedial arcuate nucleus (dmARH), whereas α1 tanycytes line the VMH and DMH. α tanycytes also contact *en-passant* different neural cells before ending on blood-brain barrier vessels and are mainly described as fuel sensors and neuronal modulators.

Unfortunately, this classification is usually too simplistic and no longer adequate regarding the recent advances in our understanding of ependyma physiology^7, 8^. Indeed, although marker genes are used to roughly separate tanycyte subtypes and typical ependymal cells, many genes exhibited a gradual ventrodorsal pattern along the 3V rather than a clear-cut distribution across subpopulations^8^. Additionally, tanycytes lining the vmARH display morphological and functional features of α or β subgroups depending on the metabolic state of the animal, revealing complex and dynamic heterogeneity^16^. Uncovering this complexity is crucial to apprehend the tanycyte’s role and functional plasticity in metabolism.

To investigate the transcriptomics behind this heterogeneity, we used fluorescence-activated cell sorting (FACS) associated with single-cell RNA sequencing (scRNAseq) to profile the typical ependymal cells’ and tanycytes’ transcriptional signatures to elucidate the dynamical changes resulting from an energy imbalance in adult mice. Our data first confirmed that our current tanycyte classification is inadequate as numerous genes overlap between subpopulations, and their heterogeneity further increases with the fasting time course. Thus, we propose pseudo-spatial and temporal analysis to elucidate these neuroanatomical and fed→fasting dynamics in gene expression along the 3V. This approach allowed us to distinguish specific *versus* overlapping markers among the ependymal subpopulations, uncover high dynamics for the subgroup facing the ARH-VMH, and propose new criteria for cell classification.

## RESULTS

### High-resolution transcriptomic profiling of ependymocytes in the mediobasal hypothalamus

To analyze the molecular signature of MBH ependymocytes, we performed FACS-associated scRNA sequencing to obtain a single-cell suspension enriched in our cells of interest (**Figure 1A**). To do so, TAT-CRE fusion protein was first stereotactically infused into the 3V of tdTomato^loxP/+^ Cre reporter mice to induce tdTomato expression in the ependyma lining the MBH (**Figure 1A**). As TAT-CRE targets cells close to the infusion site^33^, this injection allows for precisely labeling the 3V ependyma, including both typical ependymal cells and tanycytes (**Figure S1A**), as described previously^15^. One week later, MBH microdissections were harvested at 8-9 a.m. from mice under different metabolic conditions (*i.e.*, fed, 12h-fasting, and 24h-fasting) (**Figure 1A, Figure S1B**). After cell dissociation, TdTomato-positive cells were sorted by fluorescence-activated cell sorting (FACS) with a loose gating strategy to optimize the collection of ependymocytes at the risk of sorting a few neighboring cells (**Figure 1A, Figure S1C**). Our ependyma-enriched cell suspension was finally processed by chromium 10x genomics to obtain the transcriptional profile of 13 121 individual cells (**Figure 1A**, **Table S1A**).

**Figure 1.**
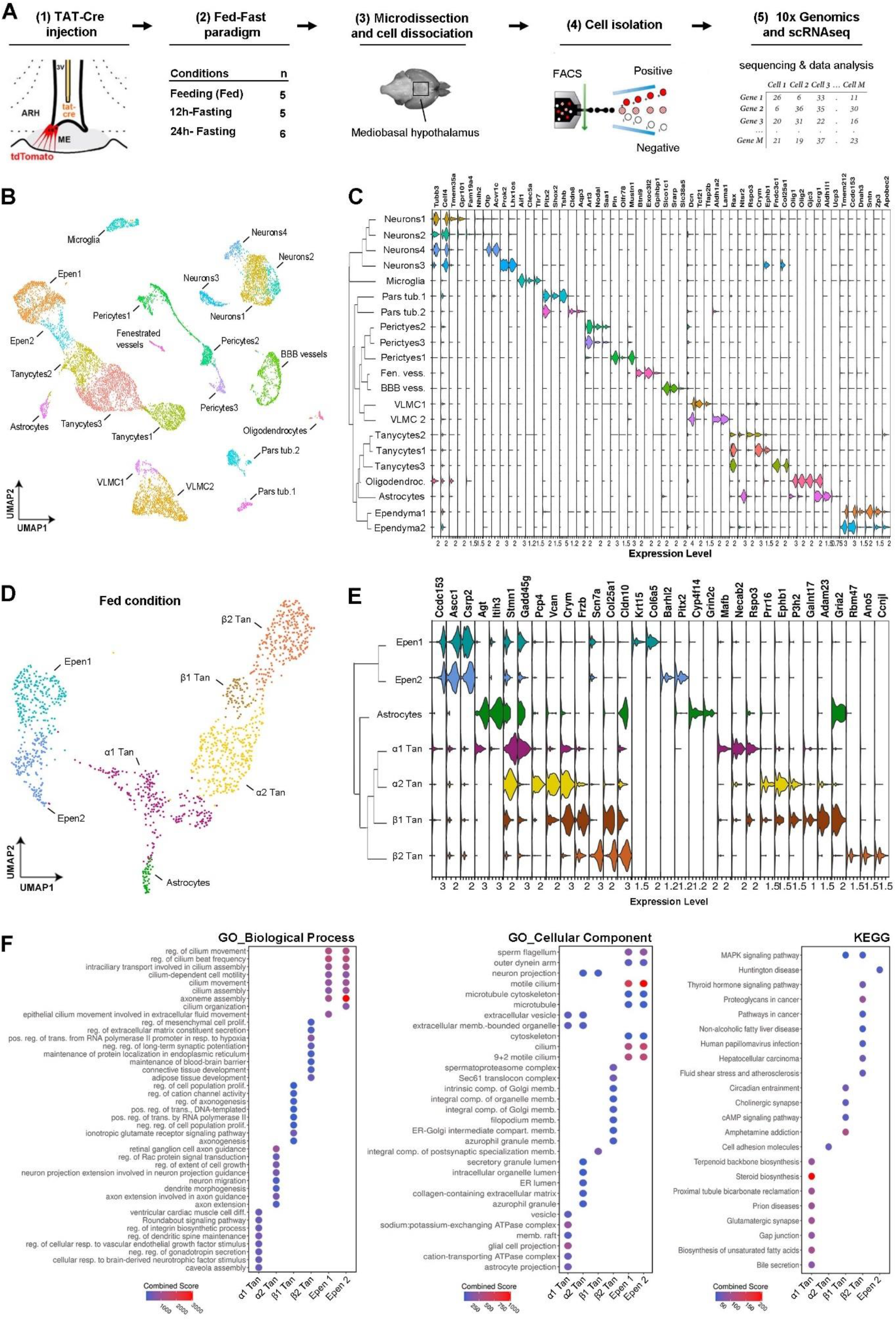
High-resolution transcriptomic profiling of MBH ependymal cells. **(A)** Experimental workflow. TAT-Cre was infused into the 3V (1) to target the ependyma. One week later, mice were sacrificed in three metabolic conditions (2). The mediobasal hypothalamus (MBH) was microdissected (3); cells were then dissociated, recovered by FACS (4), and processed by 10X 3’ whole transcriptome scRNAsequencing (5). **(B)** UMAP representation of the three integrated conditions (*i.e.*, fed, 12h-fasting, and 24h-fasting) for a total of 13’121 cells, colored per cluster and annotated according to known cell types. **(C)** Dendrogram showing the hierarchical clustering between populations and violin plot displaying known markers and specific features per cell type. Specific features are based on the ratio between the percentage of expressing cells in the given population *versus* all other cells. **(D)** Subset and re-clustering of ependymal populations in fed condition, colored per cluster and annotated according to known cell types. **(E)** Dendrogram showing relatedness between clusters and violin plot displaying enriched (left) *versus* specific (right) features per cell type. Enriched features are based on differential gene expression analysis. Specific features are based on the ratio between the percentage of expressing cells in the given population *versus* all other cells. **(F)** Cluster characterization with Gene Ontology (GO) highlighting molecular function, cellular component, and KEGG. The top 8 significant terms are displayed and classified by Enrichr combined score. BBB, blood-brain barrier; Epen, typical ependymal cells; FACS, Fluorescence Activated Cell Sorting; GO, gene ontology; Tan, tanycytes; VLMC, vascular and leptomeningeal cells.

Unsupervised clustering analysis was first performed from the UMAP embedding and identified 21 distinct subpopulations (**Figure 1B, Figure S1D-E, Table S1B**). Following the identification of their enriched features and comparing them with known marker genes, we assigned a single identity to each cluster: tanycytes (three clusters; *Rax+*), typical ependymal cells (two clusters; *Ccdc153+*, *Tmem212+*), vascular and leptomeningeal cells (VLMC) (two clusters; *Dcn+*), neurons (four clusters; *Tubb3+*), blood vessels (two clusters; *Slco1c1+* and *Exoc3l2+*), pericytes (three clusters; *Art3+*), microglia (one cluster; *Aif1+*), pars tuberalis (two clusters; *Pitx2+*), astrocyte (one cluster; *Aldh1l1+*), oligodendrocytes (one cluster; *Olig1+*) (**Figure 1C, Table S2A**). Regarding tanycyte populations, a further allocation based on our current classification may be done: “tanycytes 1” express *Crym* and *Ephb1*, corresponding to β1/α2-tanycytes; “tanycytes 2” express expressing *Rspo3* and *Ntsr2* defining α1-tanycytes; and “tanycytes 3” express *Fndc3c1* and *Col25a1* corresponding to β2-tanycytes^34, 35^ (**Figure 1C, Table S2A**).

This unsupervised clustering analysis first confirmed that our single-cell suspension was enriched in ependymocytes. Indeed, we obtained a 41.8% enrichment in both typical ependymal cells and tanycytes, the other cell populations representing less than 11% per cell type (**Table S1B**). Moreover, to confirm the specificity of our targeting approach (*i.e.*, TAT-Cre injection in the third ventricle), we analyzed the expression of tdTomato in our dataset. Consistently with our previous reports^15, 16^, our results show that TAT-Cre injection into the 3v mainly targets the ependyma: 78.3% of tdTomato-expressing cells are ependymocytes, whereas other cell populations represent less than 3% per cell type (**Figure S1E**, **Table S1C**). These different cell types likely arise from the gating strategy and/or cross-contamination of tdTomato-positive cell debris during tissue dissociation. Indeed, tanycytes make numerous contacts with various cell types in the MBH^15^, which may be kept by our soft dissociation, leading to their isolation. Finally, the hierarchical structure of the clusters revealed the impact of spatial organization and microenvironment on the cellular transcriptional profile, which appears to be organized into four categories: the parenchymal cells (*e.g.*, neurons and microglia), the pars tuberalis, the vascular system (*e.g.*, pericytes, endothelial cells, and mural cells), and the ependyma (*e.g.*, tanycytes and typical ependymal cells) (**Figure 1C**).

### Ependymocytes form a heterogeneous population along the 3V

Our FACS-associated scRNAseq approach aimed to get a high-resolution transcriptomic profile for the ependymocytes lining the MBH. To reveal finer heterogeneity along the ependyma eventually hidden in the initial clustering workflow, we next subset the data to focus on tanycytes, typical ependymal cells, and astrocytes in the fed condition (**Figure 1D, Table S1D**). After re-clustering using the same parameters, seven clusters were detected in the fed state, including two typical ependymal cell clusters, one astrocyte-like cluster, and four tanycyte clusters (**Figure 1D**). We first observed that typical ependymal cells are organized in two distinct populations, namely Epen1 (*Krt15*+/*Col6a5*+) and Epen2 (*Barhl2*+/*Pitx2*+) (**Figure 1E**). Interestingly, some features, such as *Ccdc153* or *Tmem212*, are shared with a similar expression level between these two subgroups, whereas others display different expression levels or are specific to each (**Figure 1E, Table S2B**). For instance, *Ascc1* is enriched in Epen1, whereas *Csrp2* is in Epen2 (**Figure 1E, Table S2B**). By cross-referencing each specific feature with *in situ* hybridization data from the Allen Mouse Brain Atlas^36^, no well-defined regions were found along the 3V for these two populations but rather a sparse and interlaced distribution (**Figure S2**). Next, gene enrichment analyses were performed with the enriched genes for each subpopulation to further characterize their main biological processes, cellular components, and KEGG, revealing no clear and distinct functions between these two populations. As expected, the two typical ependymal cell populations are enriched in genes related to cilium assembly and motility, consistent with their role in CSF flow regulation (**Figure 1F**). However, it is worth noticing that genes enriched in Epen1, such as *Krt15* or *Col6a5*, are involved in the extracellular matrix and related to cell communication.

Regarding the tanycyte population, known markers were first used to identify the four current subtypes, named β2-, β1-, α2-, and α1-tanycytes (**Figure 1D-E**). Using this approach, we confirmed most of the known tanycytes functions by gene enrichment analysis (**Figure 1F, Table S2B-E**). For instance, β2 tanycytes express genes related to the maintenance of the blood-brain barrier (**Figure 1F**), primarily associated with vascular development, such as *Vegfa*, and tight or adherens junctions, such as *Cldn10*, *Cldn3*, *Jam2*, and *Tjp1* (**Figure 1E, Table S2B**), validating their organization as a blood-brain barrier^16, 37, 38^. We also found genes related to the thyroid hormone signaling pathway, such as *Dio2*^21, 39^, or MAPK/ERK signaling pathway^19, 40^. Additionally, a few novel functions were also highlighted, such as extracellular matrix, endoplasmic reticulum, and Golgi compartments in β2-tanycytes. For instance, an enrichment in connective tissue growth factor (*Ctgf*) was also observed in the β2 population (**Table S2B**): this extracellular matrix-associated heparin-binding proteins notably bind different growth factors, such as VEGF^16^ or TGF-β^41^, known to be involved in ME plasticity. Regarding the transcriptomic profile of β1-, α2-, and α1-tanycytes, many genes appear to be involved in neuronal processes, especially axonal guidance, dendrite morphogenesis, and synaptic functions, highlighting tanycyte-neuron interactions (**Figure 1F**, **Table S2C-E**). Finally, confirming previous results^42^, α1 tanycytes appear to be the population closest to astrocytes, expressing genes related to astrocyte projection (**Figure 1F**).

Nonetheless, the clustering analysis also includes some limitations while analyzing the ependyma, as features may display different patterns. First, a few features are highly specific (**Figure 1E**) and characterize well-defined regions along the 3V based on *in situ* hybridization data available on the Allen Mouse Brain Atlas^36^ (**Figure S2**). Such specific features are numerous for typical ependymal cells or β2-tanycytes, which constitute stable subgroups, whereas it remains challenging to find specific markers for β1-, α2, and α1-tanycytes (**Figure 1E**). Additionally, many enriched features -calculated by differential gene expression-do not characterize one subpopulation but often span over several subgroups (**Figure 1E**), shaping ventro→dorsal or dorso→ventral gradients in gene expression along the 3V (**Figure S1H, S2**). These overlapping markers partly explain the related GOs found for the tanycyte populations, especially β1- and α2-tanycytes. Finally, some features partly characterize the subgroups, as indicated by the bimodal distribution in gene expression of some genes (*e.g.*, *Adam23* in β1-tanycytes) (**Figure 1E**). Indeed, many sparse markers are expressed by less than 80% of the cell population (**Table S2B**), suggesting finer heterogeneity among the subgroups.

Altogether, clustering analysis on the 3v ependyma fails to find clear lines of demarcation between different subpopulations, especially β1-, α-2, and α1-tanycytes, and detect the fine heterogeneity within these subgroups, raising questions regarding our current ependymal classification^8^.

### Pseudospatial analysis dissects 3V complexity

To surpass the limitation of clustering analysis regarding the characterization of the tanycyte population, we devised a pseudospatial analysis to explore the gradual pattern of features expressed along the 3V (**Figure 2A-B**). Indeed, the neuroanatomical distribution of tanycytes is crucial in establishing their identity and functions, as they project in different brain nuclei and contact various partners. Interestingly, the UMAP distribution of ependymocytes follows the anatomical ventrodorsal axis for the different ependymal subgroups (*i.e.*, β2>β1>α2>α1>typical ependymal cells) (**Figure 1B, D**). Additionally, the gradual patterns of known features, such as *Rax*, *Rarres*, or *Notch*^43^, also follow this peculiar UMAP anatomical-like distribution (**Figure S1-2**). Taking advantage of this characteristic, we used Monocle3 to sort the cells in the UMAP space that we considered as a pseudospatial trajectory from typical ependymal cells to β2 tanycytes (**Figure 2A**): this analysis allows us to reorganize and classify genes according to their pseudospatial trajectory (**Figure 2B**).

**Figure 2.**
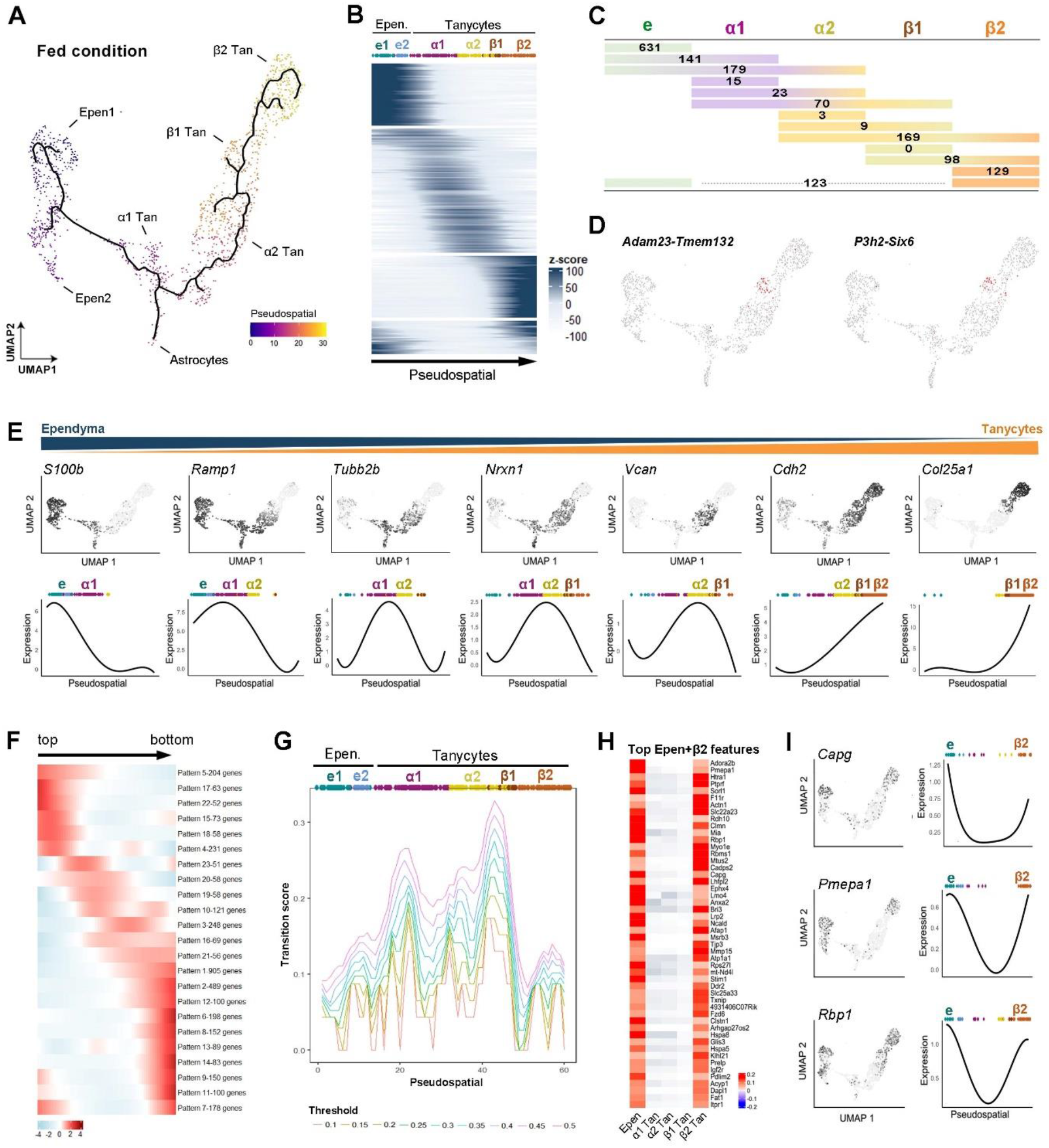
Fine heterogeneity of the 3V ependyma revealed by pseudospatial analysis. **(A)** UMAP representation displaying the pseudospatial trajectory from typical ependymal cells to β2-tanycytes**. (B)** Heatmap showing gradual gene expression in the typical ependymal cells→β2-tanycytes trajectory. **(C)** Number of features significantly correlating with one *versus* multiple ependymal populations. **(D)** UMAP representation displaying cells (in red) co-expressing *Adam23-Tmem132* and *P3h2-Six6*, characterizing the β1 population. **€** Feature plots of some overlapping genes following the typical ependymal cells→β2-tanycytes trajectory. **(F)** Heatmap showing different shared pseudospatial gene expression patterns along the top→bottom trajectory. Only twenty-three patterns with more than 50 genes are displayed. **(G)** Graph representing the main transition region along the pseudospatial trajectory. The transition score was calculated as an on-off switch in gene expression along the trajectory. Values represent on-off gating thresholds. **(H-I)** Heatmap (H) and feature plot (I) of representative genes following the U-shape pattern (*i.e.*, expression in typical ependymal cells and β2-tanycytes).

To further characterize our current classification and define relevant features, we first divided the pseudospatial trajectory by distinguishing β2-, β1-, α2-, α1-tanycytes, and typical ependymal cells based on the UMAP clustering in the fed condition. Because the two typical ependymal populations were found to share most of their features (*i.e.*, up to 70%), they were considered a single population for the rest of the analysis. We then applied a masked correlation analysis (see methods) to differentiate features highly expressed in one population (*i.e*., specific markers) from those spanning over multiple populations (*i.e.*, overlapping markers) (**Figure 2C**). Our results first identified 631 specific features (*e.g.*, *Pcp4l1*, *Ccdc153*, *Tctex1d4*) in typical ependymal cells and 129 in β2-tanycytes, confirming the robustness of these two subgroups. Based on these specific features, gene enrichment analysis for β2 tanycytes confirmed known and specific functions, such as the VEGF signaling pathway as the top KEGG term^16^. In the context of metabolism, β2 tanycyte functions include multiple terms related to lipid metabolism, such as postprandial triglyceride response (GWAS), waist circumference adjusted for BMI in smokers (GWAS), cholesterol metabolism (KEGG), regulation of lipolysis in adipocytes (KEGG) (**Table S3**). In contrast to β2 tanycytes, very few genes exclusively correlate to α1 (15 features; *e.g.*, *Fcgr2b*, *Mafb*, *Eef1a2*), α2 (3 features; *i.e.*, *Col6a3*, *Sema3a*, *Pcp4*), and β1 (0 feature) tanycytes along the pseudospatial trajectory (**Figure 2C**, **Table S3**). On the contrary, most identified features were highly correlated to multiple ependymal populations (**Figure 2C,E**). Indeed, we identified 179 features overlapping between typical ependymal cells, α1-, and α2 tanycytes and 169 features between β2-, β1-, and α2 tanycytes. Their shared functions include axon guidance and synaptic potentiation (**Table S3**). To finally validate our novel analytic approach, we applied the same workflow on the Hypomap dataset^44^: similarly, specific features were found for ependymal cells and β2 populations, whereas β1-, α2, and α1 tanycytes mainly display overlapping features (**Figure S3**).

The low number of specific markers for tanycyte subgroups, especially for β1-tanycytes, questions the pertinence of the current classification and our clustering analysis. Thus, we evaluated whether β1-tanycytes form a particular subpopulation or an overlapping mix of β2 and α2 tanycytes. To do so, combinations of features were analyzed: by combining β2/β1 and β1/α2 overlapping features, we observed that some tandems, such as *Adam23*-*Six6* or *Adam23*-*Col25a1*, appear to be specifically expressed in β1-tanycytes (**Figure 2D, Table S3**), suggesting that this population may not be stable but dynamically characterized as co-expression of overlapping markers.

Therefore, to further challenge our current classification and identify novel boundaries along the 3V only based on gene expression, we next performed an unsupervised pseudospatial pattern analysis using TradeSeq^45^. This analysis allowed us to group sets of genes following similar expression patterns along the pseudospatial trajectory: in the fed condition, twenty-three patterns containing more than 50 genes were identified (**Figure 2F**, **Figure S4**). Using these twenty-three patterns, we next detected the main area boundaries on the pseudospatial trajectory. To do so, we determined -for each pattern-transitional zones defined as regions where a switch between expression *versus* no expression occurs. (**Figure 2G**). Using this gene expression driven pseudospatial analysis, two main boundaries were found along the 3V, defining three subgroups that do not correspond to our current tanycyte subgroups (**Figure 2G**).

Finally, both supervised (**Figure 2C-E)** and unsupervised (**Figure 2F-G**) pseudospatial analysis revealed that ventrodorsal expression gradients are not the only patterns along the 3V: indeed, features can also be shared between non-neighboring populations, drawing U-shape pattern along the pseudospatial trajectory (**Figure 2H-I, Table S3**). Notably, we identified 123 features concomitantly correlating to ependymal and β2-tanycytes (**Figure 2C**, **Table S3**) and validated them using the Allen Brain Atlas (**Figure S2**). Within the genes exhibiting an Epen+β2 pseudospatial distribution, we found *Capg*, *Pmepa1*, and *Rbp1* (**Figure 2I, Table S3**), genes involved in growth factor binding, lipid binding, and endosome membrane. Globally, GO analysis revealed diverse biological processes such as hormone biosynthetic and metabolic processes, morphogenesis, and TGFB receptor signaling, among others (**Table S3**). Additionally, our pseudospatial pattern analysis also shows U-shaped patterns covering other subpopulations (**Figure 2F**).

Altogether, the pseudospatial gradient analysis challenged the current ependymal classification and highlighted the importance of considering gene expression overlaps and co-expression to characterize cell populations along the 3V more precisely. Based on specific *versus* overlapping markers, two stable populations, namely typical ependymal cells and β2-tanycytes, were identified, whereas the called α1-, α2-, and β1-tanycytes correspond to a continuum parenchymal tanycyte group with various overlapping features (**Figure 7**).

### Fasting increases cellular heterogeneity along the 3V

In addition to their heterogeneity, tanycytes are highly plastic cell types and display changes in gene expression during energy imbalance^16, 26^. To identify dynamics in gene expression profile during the fed→fasting paradigm, we subclustered the 12h-fasting and 24h-fasting datasets focusing on ependymal cells (*i.e.*, typical and tanycytes) and astrocytes and applied the same workflow used in the fed dataset condition (**Figure 3A, Table S1D**). Interestingly, more tanycyte subgroups are observed as we advance in the fasting paradigm. Indeed, five tanycyte clusters were defined in the 12h-fasting state and six in the 24h-fasting state (**Figure 3A**). Similarly, four typical ependymal cell clusters were found in the 24h-fasting *versus* only two in the fed and 12h-fasting states (**Figure 3A**).

**Figure 3.**
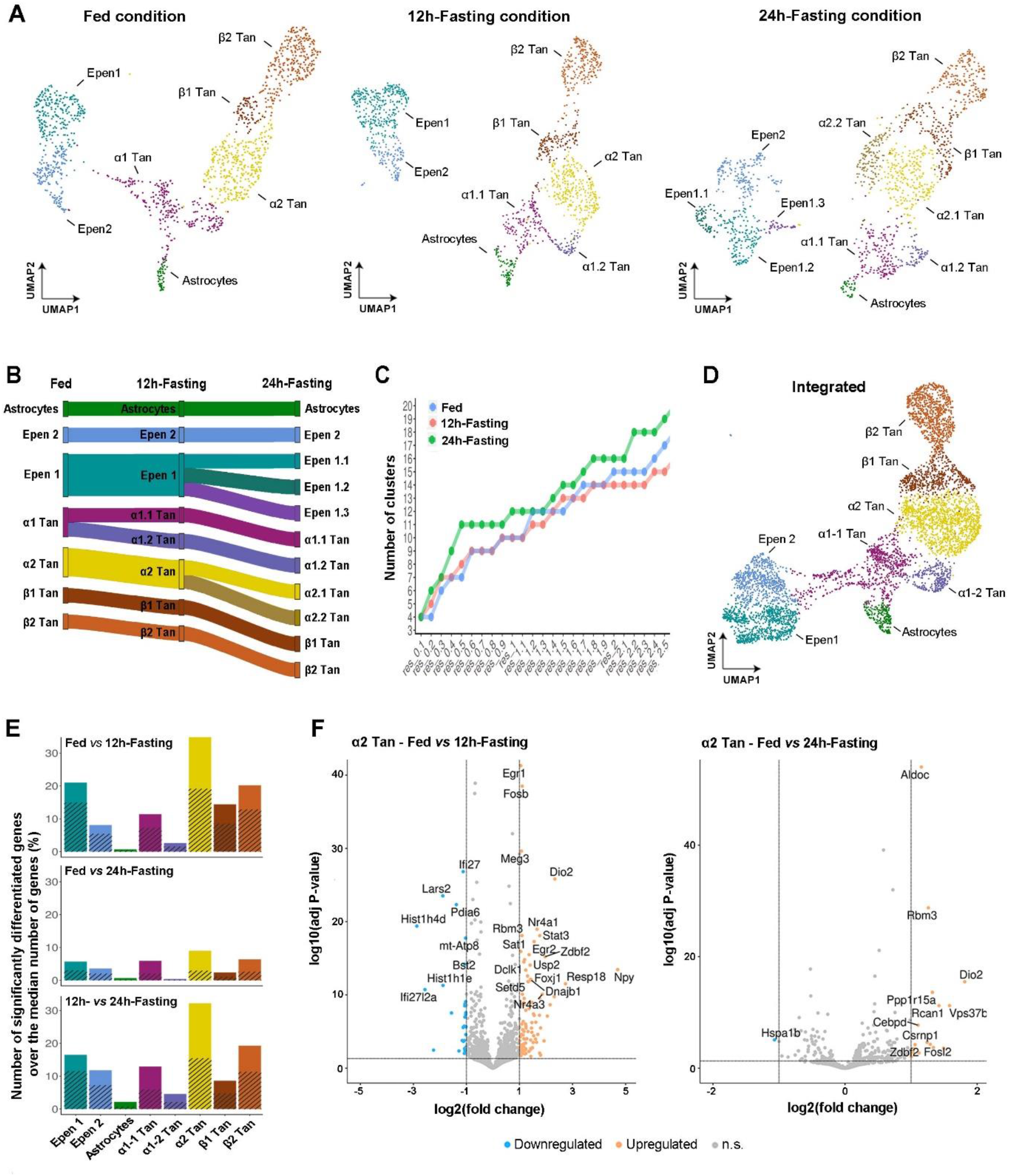
Increased heterogeneity along the 3V during energy imbalance. **(A)** Re-clustering of tanycytes, ependymal cells, and astrocytes-like cells in fed, 12h-fasted, and 24h-fasted adult male mice, colored per cluster and annotated according to known cell types. **(B)** Sankey diagram describing cell-type fate across metabolic conditions. **(C)** Number of identified clusters depending on the resolution parameter. **(D)** Integrated UMAP representation holding the three metabolic conditions together, colored per cluster and annotated according to known cell types. **(E)** Bar height representing the normalized proportion of differentiated genes calculated for each population: the number of differentially expressed genes (positive and negative) was divided by the median number of genes for each population. The shaded area of the bar represents the proportion of down-regulated genes. **(F)** Volcano plot displaying genes differentially expressed in 12h-fasting *vs.* fed and 24h-fasting *vs.* fed for the α2-tanycytes.

Based on the feature lists obtained for each subgroup in each metabolic condition (**Table S4A**), we determined the number and percentage of shared features for each combination (**Table S4B**). We concluded that β2 and β1 are stable over time, whereas α2 and α1 become more diverse (**Figure 3B**). Indeed, the α1-tanycyte population splits in two during the fed→12h-fasting transition, whereas the α2 population divides in two during the 12h-fasting→24h-fasting transition (**Figure 3B**). Regarding the split of the α1 population during 12h-fast, the α1.2 population expresses numerous ribosomal (*e.g.*, *Eef1a2*, *Rpl3*, *Rpl18*, *Rpl27a*) and mitochondrial (*e.g.*, *Atp2a2*, *Atp5b*) proteins, suggesting a high degree of translation and activation (**Table S4A**). Similarly, regarding the split of the α2 population during 24h-fasting, the α2.2 population expresses numerous ribosomal genes, whereas the α2.1 population is enriched in transcription factors (**Table S4A**). Additionally, the analysis of the shared features showed that α2.1-tanycytes in the 24h-fasting condition shared more features (*e.g.*, *Emd*, *Btg2*, or Nr4a1) with β1- and β2-tanycytes than during the fed state (**Table S4A-B**), suggesting a shift towards β-tanycyte phenotype. Many of these novel shared features are transcription factors, notably belonging to the AP1 transcriptional complex. Regarding typical ependymal cells at 24h-fasting, the epen1.3 cluster also expresses more features initially characterizing α1-tanycytes, suggesting a change toward tanycyte functions (**Table S4A-B**). To summarize, α1-, α2-, and β1-tanycytes, which were previously identified as the continuum parenchymal tanycyte group, display the most heterogenous and plastic phenotype during the fed→fasting transition.

Finally, the limitations of clustering analysis to identify tanycyte subgroups raise the question of the validity of this increased heterogeneity. To test this, we incrementally changed the resolution parameter for the clustering analysis: once again, the number of clusters increases quicker and higher at 24h-fasting compared to the other conditions (**Figure 3C**). However, the 12h-fasting heterogeneity is questionable as not always observed (**Figure 3C**).

### A functional switch occurs in tanycytes during fasting

Since tanycytes appear sensitive to the metabolic state, we next assessed differences in gene expression across the conditions. By processing ependymal cells and astrocytes from the different metabolic conditions altogether (**Figure 3D**), we detected five clusters for tanycytes, one for astrocytes, and two for ependymal cells. After establishing the identity for each group, we again revealed a greater heterogeneity for the α populations, with two subgroups for the α1 tanycytes (*i.e.*, α1-1 and α 1-2) (**Figure 3D**, **Table S4C**).

Using a pseudobulk differential analysis, we identified hundreds of genes significantly up- or down-regulated in response to fasting (**Figure 3E, Table S4D**). Surprisingly, very few transcriptional differences are observed in astrocytes, suggesting they are less affected by metabolic changes (**Figure 3E**). Conversely, α2-tanycytes appear to be the more dynamic population (**Figure 3E**). Notably, transcriptional changes in this population mainly affect RNA synthesis and processing (**Figure 3F**, **Table S4D-F**). First, numerous genes upregulated during fasting code for transcription factors, such as *Egr1*, *Egr2*, *Fosb, Nr4a1*, *Nr4a3*, *or Stat* (**Figure 3F**, **Table S4D-F**), suggesting changes in transcriptional activity. Moreover, numerous RNA binding proteins in the nucleus and the spliceosome (*e.g*., *Luc7l3*, *Rbm3*, *Srsf2*, *Srrm2*) are upregulated in 12h- or 24h-fasting (**Figure 3F**, **Table S4D-F**). Additionally, genes involved in the thyroid hormone signaling pathway and synthesis (*e.g.*, *Dio2*, *Rcan1*, *Notch2*) are also upregulated in these conditions in the α2 population (**Figure 3F**, **Table S4D-F**), consistent with previous reports^26, 39^. Regarding the other tanycyte populations (**Table S4D-F**), *Cldn10* mRNA increases in β2- and β1-tanycytes in 12h-fasting, consistent with the rise in tanycyte barrier tightness observed during fasting^16^.

Finally, while many genes are differentially regulated in specific subtypes, several genes changed similarly across the different tanycyte populations. For instance, between fed and 12h-fasting conditions, 294 and 474 genes are significantly upregulated (FDR<0.05) in β2- and α2-tanycyte subgroups, respectively, with 131 shared genes (**Table S4E**). Notably, aldolase (Aldoc), an enzyme involved in glycolysis/gluconeogenesis, is the most upregulated gene in all cell types in the 24h-fasting state (**Figure 3F**, **Table S4D**), suggesting changes in metabolic pathways. Similarly, *Mt1* and *Mt2* genes, encoding metallothionein-1 and 2, are upregulated in all populations after 24h-fasting (**Table S4D**). These low molecular weight proteins rich in cysteine bind divalent heavy metal ions and play several putative roles in metal detoxification, Zn and Cu homeostasis, and scavenging free radicals. Interestingly, disrupting these two metallothionein genes in mice resulted in obesity^46^.

During energy unbalance, the temporal factor plays a crucial role: different processes are activated at 12h-fasting *versus* 24h-fasting. Three time-points (*e.g.*, fed, 12h-fasting, and 24h-fasting) being used in our experimental approach, we next analyzed gene expression dynamics in tanycyte subpopulations through the fed→fasting temporal trajectory (**Figure 4**). To do so, we performed a stepwise differential expression analysis to classify genes into eighteen discrete temporal patterns (**Figure 4A-B**, **Table S5**) characterized as “increasing”, “transitional”, and “decreasing” trajectories. The expression of “increasing” genes significantly increases in 24h-fasting *versus* fed state while taking different routes at 12h-fasting (**Figure 4A-B**). Conversely, gene expression in the “decreasing” group significantly decreases in 24h-fasting *versus* fed state while taking different routes at 12h-fasting (**Figure 4A-B**). For the “transitional” trajectory, no significant differences were found between fed and 24h-fasting, while gene expression significantly increases or decreases at 12h-fasting (**Figure 4A-B**). We next investigated Gene Ontology terms (GO) in single or combined trajectories (**Figure 4C, Table S5**). For the α2-population, genes related to asymmetric synapse and autolysosome increased with fasting (*i.e.*, I1+I4 or I1+I2+I3+I4 combinations). Concomitantly, genes involved in ribosomal assembly and translation are upregulated at 24h-fasting. Conversely, endoplasmic reticulum function, protein processing, and vesicular functions are downregulated in fasting (*i.e.*, D1+D4 or D1+D2+D3+D4 combination), suggesting that protein processing is replaced by RNA processing in fasting. Interestingly, the transitional temporal trajectories (*i.e.*, I6+T6+D6 combination for an upregulation and I5+T5+D5 combination for a downregulation at 12h-fasting, respectively) indicate a stepwise functional reorganization: genes involved in mRNA processing, especially RNA splicing, are upregulated at 12h-fasting, while those engaged in stability and translation increase at 24h-fasting (**Figure 4C**).

**Figure 4.**
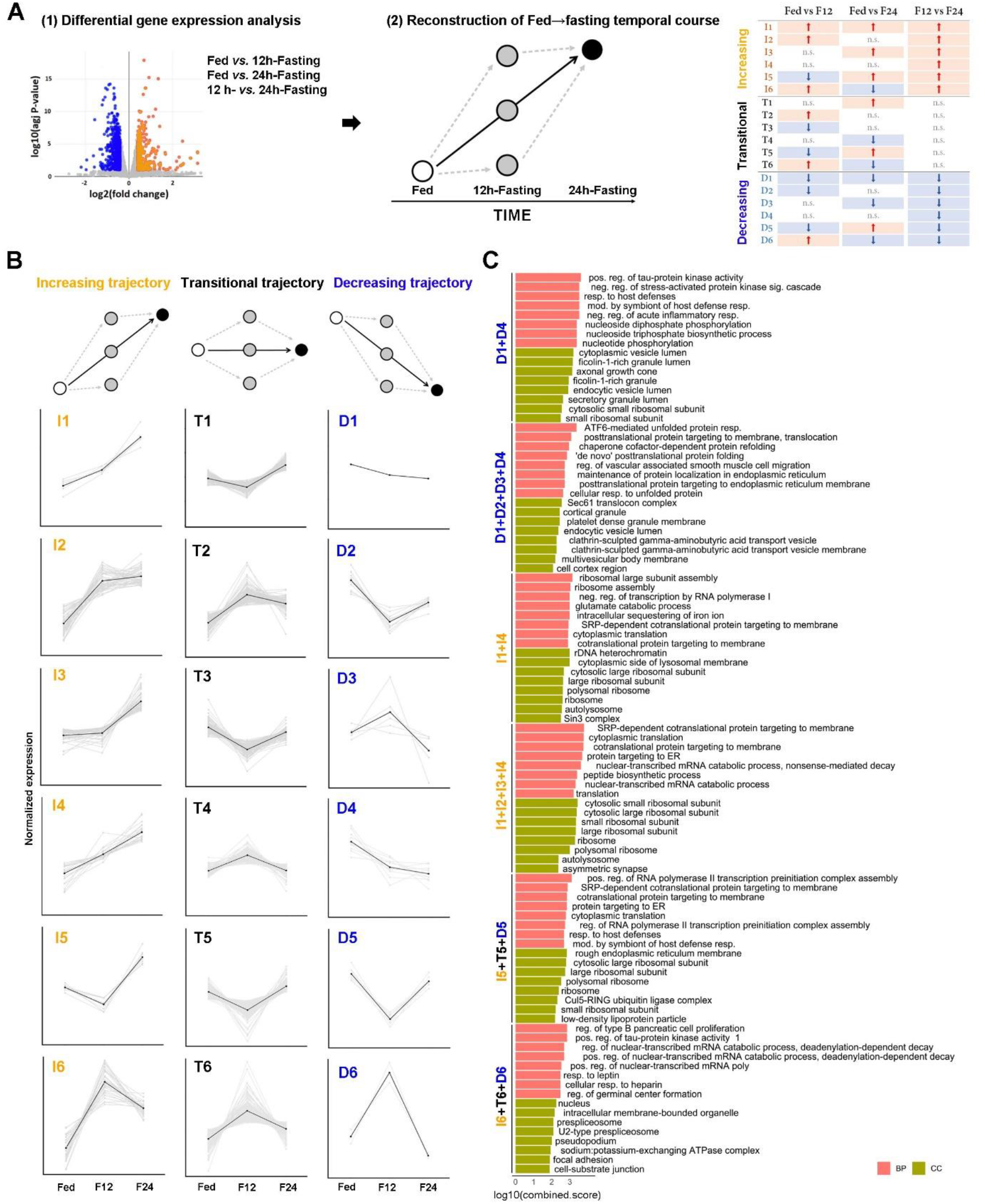
Fed→Fasting temporal trajectories based on differential gene expression reveal a cell state switch during energy imbalance. **(A)** Summary of the analytic workflow. Differentially expressed genes for each comparison (1) were integrated to draw fed→fasting temporal transitions (2) and defined eighteen trajectories (3). **(B)** Eighteen different trajectories observed for the fed→fasting temporal transitions in α2 tanycytes. **(C)** Trajectory characterization with Gene Ontology (GO) highlighting molecular function and cellular component in α2 tanycytes. The top 8 terms are displayed and classified by Combined Score. I1+I4 or D1+D4 trajectories were combined to reflect a continuous increase or decrease in gene expression, respectively. I1+I2+I3+I4 or D1+D2+D3+D4 trajectories were combined to reflect a global increase or decrease over time, respectively. I6+T6+D6 or I5+T5+D5 trajectories were combined to reflect an increase or decrease at 12h-fasting, respectively.

### Fasting redistributes gene expression gradients along the 3V

As described in our pseudospatial analysis in the feeding condition, a tanycyte classification based on clustering marker features is not entirely appropriate for the dynamic modularity of tanycytes (**Figure 2, 4**). To study the transcriptional dynamics in gene expression induced by the fed→fasting paradigm, we next performed a differential pseudospatial gradient analysis by processing the datasets from the different metabolic conditions altogether (**Figure 5**). As described previously, the pseudospatial trajectory follows the neuroanatomy from typical ependymal cells to β2 tanycytes, and cells for different feeding and fasting conditions are sparsely distributed, indicating the transcriptional identities of these trajectories are stable across those experimental conditions.

**Figure 5.**
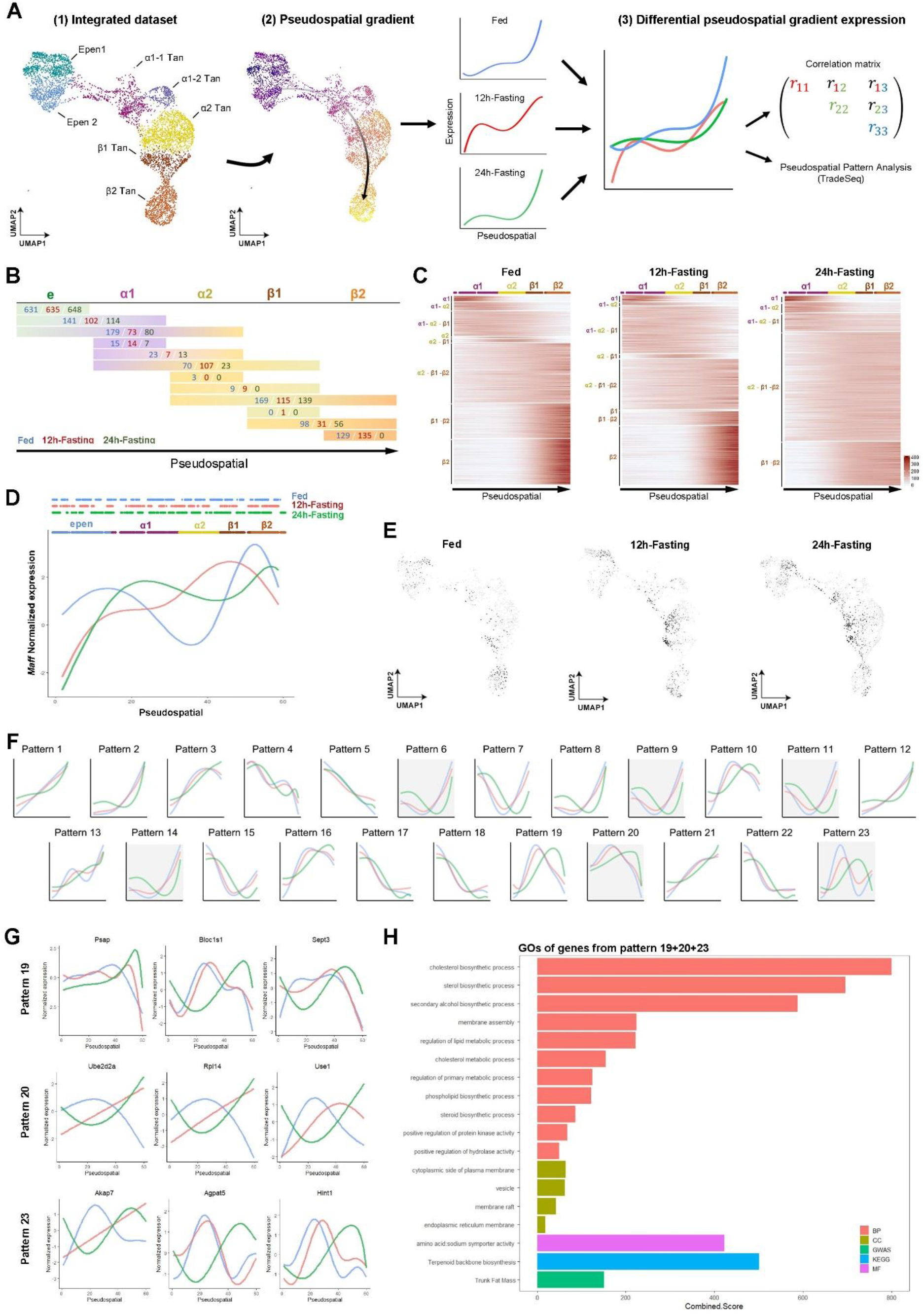
Differential pseudospatial analysis reveals a phenotype switch from α2- and β1- to β2-tanycytes during energy imbalance. **(A)** Summary of the analytic workflow. After defining the pseudospatial trajectory from typical ependymal cells to β2-tanycytes in the integrated dataset (1), the difference in the pseudospatial gradient expression between conditions was calculated either by correlation or TradeSeq (2). **(B)** Number of features significantly correlating with one *versus* multiple ependymal populations for each metabolic condition. Fasting decreases the number of specific features. **(C)** Heatmaps showing the specific and overlapping features found in the α1→β2-tanycytes trajectory. **(D-E)** Pseudospatial trajectories (D) and UMAP (E) of *Maff* in fed, 12h-fasting, and 24h-fasting conditions. **(F)** Main patterns in gene expression along the pseudospatial trajectory in fed, 12h-fasting, and 24h-fasting conditions. Patterns found in fed condition are used as reference. **(G)** Feature plots of some genes found in patterns 19, 20, and 23. **(H)** Pattern characterization with Gene Ontology (GO) highlighting biological process (BP), molecular function (MF), cellular component (CC), KEGG, and GWAS. The top terms are displayed and classified by Enrichr combined score.

By first keeping our current classification (*i.e.*, β *versus* α tanycytes), we identified overlapping and specific markers along the pseudospatial trajectory by gene expression localization across the different metabolic conditions. This analysis first revealed that, during 24h-fasting, many genes are classified as overlapping features (**Figure 5B-C**). Strikingly, no specific markers are found for the β2 population anymore. In contrast, numerous specific markers in the fed condition become β1-β2 overlapping features at 24h-fasting (around 20%), suggesting an extension of β2 phenotype to the β1 subgroup. For example, *Maff* (*i.e.*, a transcription factor involved in cellular stress response) is highly expressed in β2 populations in the fed condition, whereas it displays a shift toward α subgroups during fasting (**Figure 5D-E**).

We next analyzed the different gene expression patterns along the 3V using TradeSeq to further challenge our current classification. First, applying TradeSeq to each condition revealed an increase in the number of gene expression patterns. Indeed, while the fed state displays 23 patterns, 32 and 34 patterns were found for the 12h-fasting and 24h-fasting conditions, respectively (**Figure S4B**), suggesting an increased heterogeneity during fasting. When analyzing the switches in gene expression along the 3V, a single transition boundary was calculated at 24h-fasting (**Figure S4C**), showing that the high gene expression dynamics at this stage prevent the tanycyte classification. Additionally, we analyzed the distribution of the 23 fed patterns at 12h- and 24h-fasting (**Figure 2F**). While the fed and 12h-fasting conditions display similar gene expression distribution along the 3V, 24h-fasting induces changes in numerous patterns, notably ventro→dorsal (*i.e.*, patterns 6, 8, 9, 11, and 14) and dorso→ventral shifts (*i.e.*, patterns 19, 20, and 23) (**Figure 2F**). Focusing on patterns 19, 20, and 23, which present substantial changes, the top genes showing a shift code for proteins related to lysosomal compartments (*i.e.*, *Psap*, *Bloc1s1, Ube2d2a, Use1*), Ca2+ signaling (*i.e.*, *Akap7*, *Hint1*), and *de novo* phospholipid biosynthesis (*i.e.*, Agpat5) (**Figure 2G**). Gene enrichment analysis revealed changes in cholesterol and lipid metabolism (**Figure 2H**).

### Inferring transcriptional activity and tanycyte-neighboring cell interactions explain pseudospatial shifts in the tanycyte population

We finally attempted to explain the shifts in tanycyte phenotypes observed along the ventrodorsal pseudospatial trajectory, especially for the parenchymal tanycyte subgroup, by analyzing active regulons and cell-cell communication according to the metabolic conditions. Given the significant upregulation of TFs observed during fasting (*i.e.*, *Egr1*, *Egr2*, *Fosb*, *Nr4a1*, *Nr4a3*, or *Stat*) (**Figure 3F**), transcriptional activity is likely increased and may induce a transcriptional rearrangement. To further investigate the activity of transcription factors, we performed a SCENIC analysis^47^ in the three metabolic conditions on the pseudospatial trajectory. First, we observed that the number of active regulons detected by the algorithm dramatically increased from fed to fasting (85 in fed *vs.* 424 in 12h-fasting *vs.* 427 in 24h-fasting), highlighting once again a dynamic gene expression in these conditions. The activity of the regulons shared among all three states (59), represented as a pseudospace ordered heatmap (**Figure 6A**), shows three main distinct active regions in the fed condition, defined as the ependymal cells, a central region encompassing α1, α2, and β1, and the β2 population and confirming our reclassification (**Figure 7**). Strikingly, at 12h-fasting, many TFs, including the above-mentioned *Egr1*, *Egr2*, and *Fosb*, change their activity region from β2 to β1-α2-α1 while the ependymal specific transcriptional block stays stable (**Figure 6B**). At 24h-fasting, we notice 4 main activity regions with fewer TFs presenting with a particular ependymal and β2 expression profile, while most TFs have a broad activation profile across many cells (**Figure 6C**). This analysis confirms the increase in transcriptional activity and the transcriptional rearrangement induced by the metabolic changes, leading to a more complex and dynamic tanycyte population.

**Figure 6.**
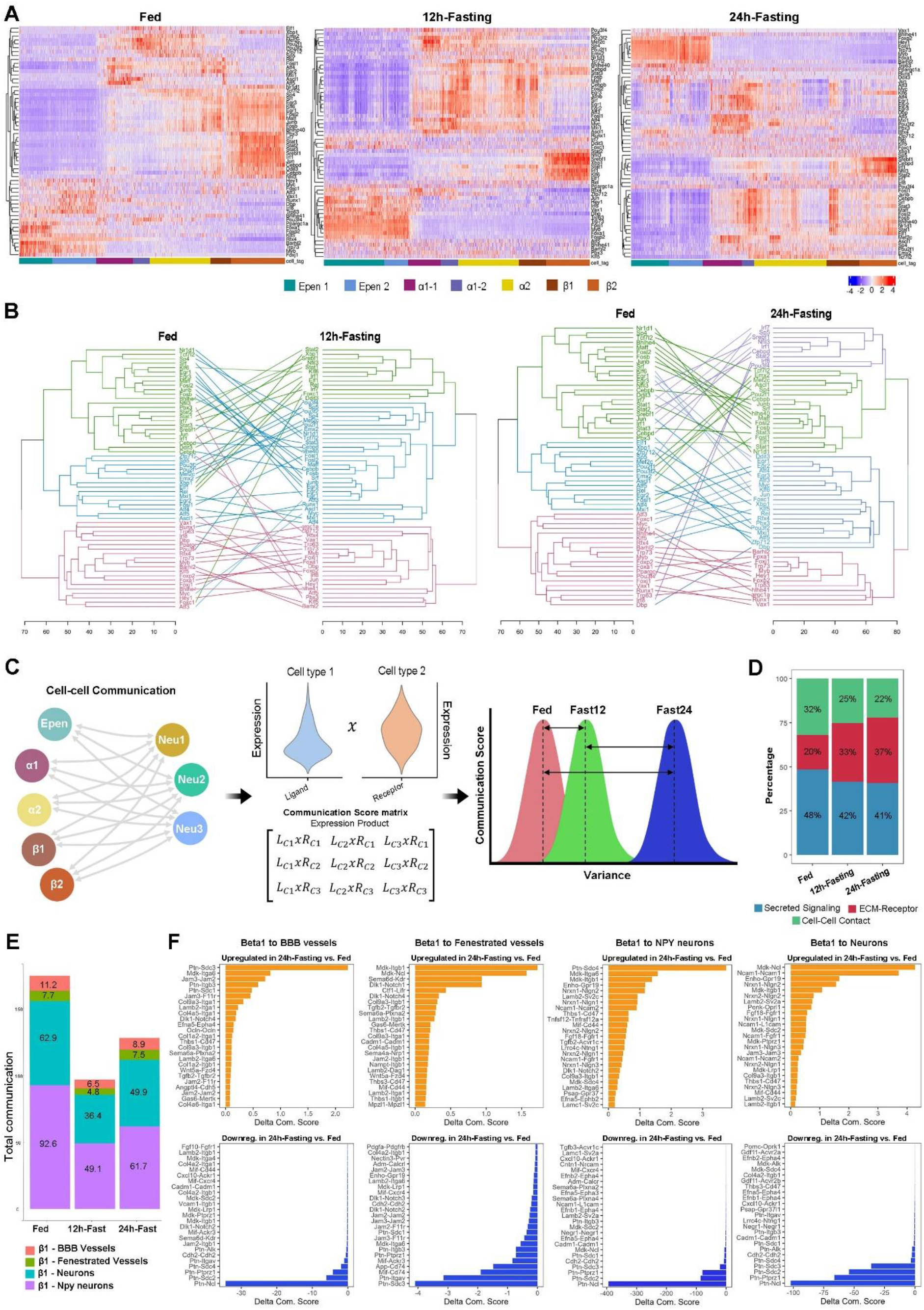
Inference of active regulons and tanycyte-neighboring cell communication for the specialization and shift of transcriptomic profile. **(A)** Heatmaps of shared transcription factor activity in the fed, 12h-fasting, and 24h-fasting conditions ordered according to the pseudospatial trajectory. **(B-C)** Tanglegrams showing the shift in transcriptional activity from the fed to 12h-fasting and the fed to 12h-fasting conditions. The first branching highlights three main clusters in fed and 12h-fasting, mostly actives in ependymal, α1-α2-β1 and β2 regions, while four clusters are derived from the first branching in 24h-fasting. Lines connecting dendrograms’ leaves across the metabolic conditions uncover many regulons changing their region of activity from β2 to α1-α2-β1. **(D)** Summary of the analytic workflow for cell-cell communication. **(E)** Proportion of each interaction type according to the metabolic condition. **(F)** Proportion of interactions with the different partners. **(G)** Histograms showing the top up and downregulated interactions with β1 populations.

**Figure 7.**
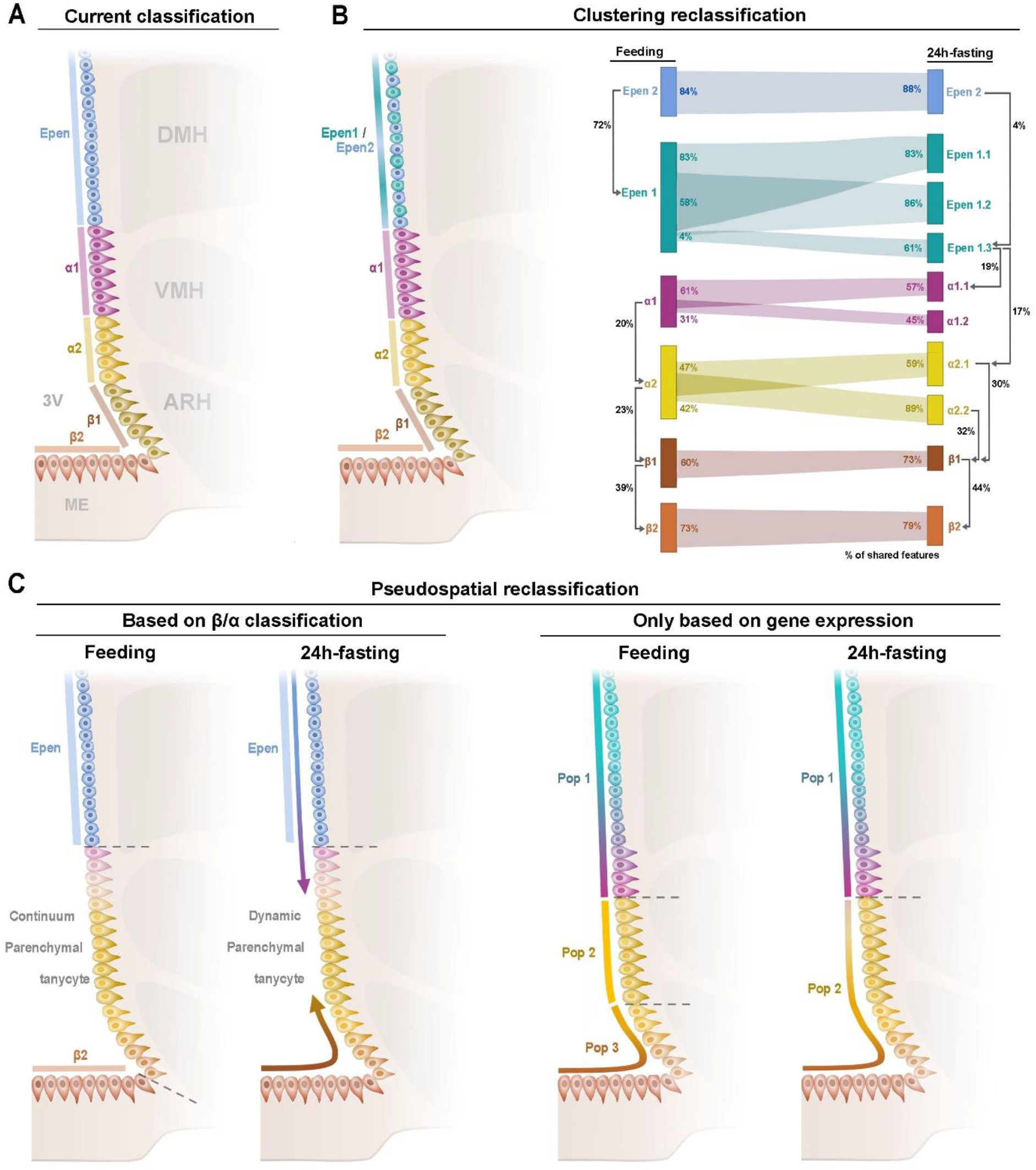
An improved model for tanycyte classification based on clustering and pseudospatial analysis. **(A)** Current ependymal classification indicates four tanycyte subgroups (*i.e*., α1, α2, β1, and β2) and one typical ependymal population. **(B)** Clustering analysis showed four tanycyte subgroups and two typical ependymal populations. During fasting, this heterogeneity increases, especially for parenchymal tanycytes and typical ependymal cells. Numerous features are shared between subgroups, and parenchymal tanycytes acquire β features during fasting (*legends for percentages* = the population at the beginning of the arrow possesses n% of shared features with the population at the end of the arrow). **(C)** Pseudospatial analysis described β2 tanycytes and typical ependymal cells as stable populations and a continuum parenchymal tanycyte group with gradual changes in gene expression. During fasting, β2 tanycyte specificity is lost, and many features are shared with other tanycytes.

Finally, tanycyte interactions with different cell types may also explain tanycyte plasticity observed during fasting. Indeed, cells sense the presence of potential interaction partners through a wide range of receptors and, specifically, respond by changing the expression of many target genes via complex regulatory networks. To study tanycyte-neighboring cell interactions, we generated a communication score matrix based on ligand-receptor expression products between tanycytes and other cell types (**Figure 6C**). Using ANOVA, our analysis allowed us to determine the dynamics of cell-cell communication between fed and fasting conditions for each pair of cell populations. As β1-α2 tanycyte populations constitute a transition subgroup with high dynamics during fasting (**Figure 2-3**), we further investigated the interactions of these cells with BBB vessels, fenestrated vessels, NPY neurons, and other neuronal populations (**Figure 6C**). First, fasting conditions increase ECM-receptor interactions and decrease communications related to secreted signaling and cell-cell contacts (**Figure 6D**). Similarly, analyzing the global communication score between β1 tanycytes and their partners revealed decreased communication during fasting (**Figure 6E**). However, the normalized communication score indicates that β1 tanycytes mainly decrease their interaction with BBB vessels while keeping a stable interaction with fenestrated vessels at 24h-fasting (**Figure 6E**), consistent with previous reports^38^. Intriguingly, the β1 tanycyte/NPY neuron interaction score decreases during fasting (**Figure 6E**). Second, regarding ligand-receptor couples between β1 tanycytes and their partners that are down- and upregulated during 24h-fasting, tanycytic pleiotrophin (*Ptn*) appears crucial to neural populations through its different receptors. Indeed, *Ptn-Sdc4* increases between β1 tanycytes and NPY-neurons during fasting, whereas Ptn-Ncl, Ptn-Ptprz1, and *Ptn-Sdc2* decrease (**Figure 6E**). Additionally, while *Ptn-Sdc4* increases with NPY-neurons, it decreases with other neuronal populations. Pleiotrophin is a secreted heparin-binding growth factor involved in cell growth and survival, cell migration, and angiogenesis. Different functions may occur through its different receptors. In particular, *Sdc4* plays a role in adiposity and metabolic complications^48^.

## Discussion

By using FACS-associated single-cell RNA sequencing, this study deciphers the heterogeneity and plasticity of the 3V to better understand its roles in regulating energy balance. Focusing on adult mice, when the 3V is well-differentiated, we proposed a novel analytic approach, namely pseudospatial trajectory analysis, to characterize cell populations according to ventrodorsal patterns in gene expression along the ventricle. In this way, we first highlighted specific markers which characterize peculiar ependymal cell subgroups *versus* overlapping features which overlap in different populations. Taking the reasoning further, we delimited dynamic regions along the 3V and compared the pseudospatial molecular patterns during a fed→fasting time course, confirming that some features can shift from one ependymal subtype to another.

For a decade, the main characteristic given to the MBH ependyma is its high heterogeneity, conferring its multifunctional biological properties. Historically, our current classification divides the 3V into five subtypes: typical ependymal cell, α1-, α2-, β1-, and β2-tanycytes. During brain development, the separation in ependymal cells *versus* tanycytes appears starting E13 in mice^49, 50^, and the differentiation in specific tanycyte subgroups is present starting P4^42, 50, 51^. Recent studies using scRNAseq and focusing on the hypothalamus were able to detect these different subpopulations and give us their molecular markers^34, 35, 44, 52^. Compared to these other datasets, our study aimed to reach a higher resolution of the MBH 3V in adult mice to further detail tanycyte classification. To this purpose, we adapted FACS associated with single-cell RNA sequencing, primarily used to study the developmental brain^42, 53^, to sort tdTomato-positive ependymal cells in adult mice in different metabolic conditions. Using a basic clustering analysis in the fed state, we also detected the four historical tanycyte subgroups with a comprehensive marker list for each and highlighted known and novel tanycyte functions. First, our results did not suggest neurogenesis from tanycytes in the adult brain. During the postnatal ages, neurons and glia arise from tanycytes and integrate hypothalamic circuits involved in the regulation of energy balance^28, 29^. Yoo *et al.* notably observed a small fraction of proliferative tanycytes at P8, from which arise neurons, oligodendrocyte progenitor cells, astrocytes, and ependymal cells, whereas, starting P17, neuronal and glial generation decreases^42^. While genetic manipulations^42^ or neural injury^54^ can reactivate tanycytes’ proliferative and neurogenic capacity, metabolic changes do not seem to impact this tanycyte function. However, a peculiar trajectory toward astrocytes was observed in our dataset, as described by others^42^, suggesting potential astrogenesis. Finally, parenchymal tanycytes (*i.e.*, α2- and β1-populations) expressed numerous genes related to synaptic structure and functions. Among these genes, many belong to ligand-receptor couples for tanycyte-neuron communication. This result is supported by our recent data showing that tanycytes contact neurons and synapses in the MBH^15^ and receive synaptoid contacts along their processes^15^. Such molecular contacts also suggest that tanycytes function as astrocytes in tripartite synapses and can modulate neuronal firing activity as previously described^25, 27^. Additionally, given that tanycytes primarily express post-synaptic molecules, they would more likely interact with presynaptic parts and modulate neurotransmitter release. Further exciting studies are necessary for decoding such tanycyte-presynaptic terminal communications.

The clustering analysis of the 3V ependyma also revealed many limitations. Many features do not well-characterize specific subgroups, either because they overlap many subgroups or display a sparse distribution within the same population. Altogether, these weaknesses highlight an arbitrariness and oversimplification of the current tanycyte classification.

While scRNA-seq is a resolutive plus value to characterize cells from complex tissues, it also leads to a loss of spatial information. However, the spatial characteristic in complex tissues such as the brain likely plays a crucial role in microdomain functions. Interestingly, the distribution of ependymal cells in our UMAP dataset follows the neuroanatomical distribution along the ventrodorsal axis from β2 tanycytes to typical ependymal cells. Adapting the pseudotime analysis to utilize the spatial information in the UMAP representation, we showed that pseudospatial analysis could accurately identify spatially relevant tanycyte subpopulations and their dynamics based on gene expression. First, our approach based on anatomically-relevant patterns allows the further characterization of our current classification. Considering specific *versus* overlapping markers and co-expressed features (for β1 tanycytes) lead to a more accurate distinction between the known subgroups along the 3v. Indeed, many available tanycyte markers actually match the overlapping features. Furthermore, delimiting ependymal regions by uniquely considering the gene expression pattern challenged our current classification, splitting the trajectory at different areas. Certainly, three main subgroups may be done along the 3V: two stable populations, namely typical ependymal cells and β2-tanycytes, with numerous specific markers and functions and a continuum parenchymal tanycyte group (corresponding to the current α1-, α2-, and β1- tanycytes) which constitute the most dynamic population with various overlapping features.

Energy unbalance affects tanycytes’ gene expression and functions^8, 16, 26^. However, most of these studies were performed at the tissue level, or at best, on FACS-isolated tanycytes, limiting the characterization of gene expression dynamics in the different tanycyte subgroups. In the literature, β2- and β1-tanycytes are mainly described for their role in regulating food intake through blood-brain exchangers. Surprisingly, our dataset revealed higher responsiveness during fasting for α tanycyte populations. This disparity with the literature may rely on the fact that many of these studies focused on the ventromedial arcuate nucleus and the median eminence, where β tanycytes are located. While few studies characterize α tanycyte populations, especially related to their glucose sensing and neuronal modulator properties, it would be interesting to define their different roles more precisely. Moreover, differential gene expression on clusters revealed a functional reorganization in tanycyte populations during fasting. More specifically, protein processing shuts down, whereas mRNA processing increases. These changes in mRNA metabolism are associated with an increased proportion of spliced mRNA, suggesting that RNA processing is crucial during fasting. Interestingly, genes related to spliceosome were also upregulated at 12h fasting. This increase in splicing could be related to mRNA stabilization to maintain some expression or to allow rapid protein translation during refeeding.

Additionally, tanycytes may switch from one subgroup to another during energy imbalance. As reported in the literature, dorsal β1-tanycytes display organized tight junction proteins following 24h-fasting and contact newly fenestrated vessels in the vmARH, resembling β2 tanycytes^16^. Our differential pseudospatial gradient analysis confirmed such a shift, especially for the continuum parenchymal tanycyte group, suggesting that a dynamic classification would better reflect tanycyte heterogeneity. In particular, our data revealed substantial shifts in genes related to lipid and cholesterol metabolism. So far, very few studies have addressed the role of tanycytes in lipid incorporation and response, but this promising field is developing. In particular, β tanycytes contain lipid droplets^12^ and metabolize palmitate to maintain body lipid homeostasis^55^. Interestingly, obese mice present alterations in tanycyte lipid metabolism and lipid droplet contents^56^. Because lipids act as signaling molecules and can promote inflammation under high fat diet^57, 58^, more studies are crucial to elucidate their role in tanycyte functions.

Altogether, revisiting and multiplying scRNAseq analytic approaches allowed us to define the heterogeneity and plasticity of tanycyte populations. The classification in α *versus* β tanycytes is not so evident while considering gene expression. Many factors, including changes in transcription factor activity and cell-cell communications, contribute to the reorganization of tanycyte gene expression through different metabolic conditions, urging us to apprehend the tanycyte population as a whole, complex, and changing continuum.

## Abbreviations

ARH: arcuate nucleus
CSF: cerebrospinal fluid
DMH: dorsomedial nucleus
FACS: fluorescence-activated cell sorting
GO: gene ontology
GWAS: genome-wide association studies
KEGG: Kyoto encyclopedia of genes and genomes
MBH: mediobasal hypothalamus
ME: median eminence
scRNAseq: single-cell RNA sequencing
UMAP: uniform manifold approximation and projection
vmARH: ventromedial arcuate nucleus
VMH: ventromedial nucleus
3V: third ventricle

## Acknowledgment

Encapsulation, library preparation, and sequencing were performed at the Lausanne Genomic Technologies Facility, University of Lausanne, Switzerland (https://www.unil.ch/gtf/en/home.html). The authors thank the CIG mouse facility (UNIL) and the mouse Metabolic Evaluation Facility (MEF-UNIL). This work was supported by the Swiss National Science Foundation (PZ00P3_174120) and European Research Council Starting Grant (TANGO, No. 948196). M.B. and F.S. are supported by the Swiss National Science Foundation (310030_185292), Horizon2020 (847941), and Novartis Foundation for medical-biological research (18A052). A.M. is supported by the Swiss National Science Foundation (310030_205068). F.L. is supported by European Research Council Starting Grant (TANGO, No. 948196), Novartis Foundation for medical-biological research, and the Swiss National Science Foundation (PCEFP3_194551). BT received support from European Research Council Advanced Grant (INTEGRATE, No. 694798) and the Swiss National Science Foundation (310030_182496).

## Author contributions

F.L. designed and performed experiments. F.L. and F.S. oversaw the research. F.S. coordinated the bioinformatic analysis. M.B. and D.L.R. analyzed the single-cell RNAseq dataset and produced the figures. F.L. wrote the manuscript. All authors edited the manuscript.

## METHODS

### Mice

2-to-4-months old male Rosa26-floxed stop tdTomato mice and C57Bl6/J mice (initially obtained from Charles River) were used in this study. Animals were housed in groups (from 2 to 5 mice per cage) and maintained in a temperature-controlled room (at 22−23°C) on a 12 h light/dark cycle with ad libitum access to a chow diet (Diet 3436; Provimi Kliba AG, Kaiseraugst, Switzerland). For genotyping, biopsies were collected, DNA extraction was performed using the HOTSHOT method, and PCR amplification was performed using KAPA2G ReadyMix Kit (Sigma Aldrich, KK5103) following the manufacturer’s instructions. Primers used for PCR are available from the authors. All animal procedures were approved by the Veterinary Office of Canton de Vaud and performed at the University of Lausanne.

### TdTomato expression in tanycytes

To induce tdTomato expression in tanycytes, TAT-CRE fusion protein (Excellgen, EG-1001) was stereotactically infused into the 3V (500nl to 2ul over 3 min at 2 mg/ml; at the coordinates form the bregma of AP= −1.7 mm; ML= 0 mm; DV= –5.3 mm (from cortex surface)) of ketamine/xylazine-anesthetized mice (100 mg/kg and 20 mg/kg, respectively) one week before experiments. The quality of the injections was arbitrarily estimated by visualizing the backflow of a CSF drop after the needle removal (**Table S1A**).

### Tissue dissociation and cell collection for single-cell RNA sequencing

Experiments were performed in fed (n=5 mice), 12h-fasting (n=5 mice), and 24h-fasting (n=6 mice) conditions one week following TdTomato induction in the ependymal layer (**Table S1A**). Mice were killed between 8 a.m. and 9 a.m. Adult male mediobasal hypothalami were microdissected using a binocular microscope and put in 500µl ice-cold papain solution (20u/ml). Cells were then dissociated following Worthington’s instructions. Briefly, cells were dissociated by incubating papain solution containing microdissected tissue at 37°C for 30 minutes, followed by gentle manual trituration. Cell suspensions were centrifuged at 330g for 5 minutes, and the cell pellet was then resuspended in a 500 µl albumin/ovomucoid protease inhibitor solution (1 mg/ml). A discontinuous density gradient centrifugation was performed by layering the cell suspension on an 800 µl albumin/ovomucoid protease inhibitor solution (10 mg/ml) and centrifuged at 100g for 7 minutes. Cell pellets were finally resuspended in 400 µl ice-cold calcium-free and magnesium-free HBSS.

TdTomato-positive singlet cells were sorted using a Beckman Coulter Moflo Astrios FAC-sorter according to their Forward and Side scattering properties (FSC and SSC), their negativity for DAPI (Viability dye, Blue DNA intercalating agent, ThermoFisher, cat nb D1306, λEx/λEm (with DNA) = 358/461 nm), their positivity for RedDot1 (Viability dye, Far-red DNA intercalating agent, Biotium, #40060-1, λEx/λEm (with DNA) = 662/694 nm), and their level of tdTomato fluorescence emission (λEx/λEm = 554/581 nm). Gating parameters were set to improve the purity (*i.e.,* fast collection, Noozle 70, purity) while keeping a loose FACS gate. Cells from identical conditions were collected and pooled in 10 ul in calcium-free and magnesium-free HBSS (Merck, H8264) + 50% FBS (Gibco, 16030074), yielding a 500-600 cells/µl suspension (**Table S1A**).

### Single-cell capturhe, cDNA library preparation, and sequencing

Sorted cells were accurately counted using a hemocytometer with Trypan blue staining to validate the viability (at least 80%). A Chromium Next GEM Chip B (10X Genomics) was loaded with approximately 5’000 cells and sequencing libraries prepared strictly following the manufacturer’s recommendations (manual CG000183 revA). Briefly, an emulsion encapsulating single cells, reverse transcription reagents, and cell barcoding oligonucleotides were generated. After the reverse transcription step, the emulsion was broken, and double-stranded cDNA was generated and amplified for 12 cycles in a bulk reaction. This cDNA was fragmented, a P7 sequencing adaptor-ligated, and a 3’ gene expression library generated by PCR amplification for 14 cycles.

Libraries were quantified using a fluorimetric method, and their quality was assessed on a Fragment Analyzer (Agilent Technologies). Cluster generation was performed with 140 pM of an equimolar pool from the resulting libraries using the Illumina HiSeq 3000/4000 PE Cluster Kit reagents. Sequencing was performed on the Illumina HiSeq 4000 using HiSeq 3000/4000 SBS kit reagents. Sequencing data were demultiplexed using the bcl2fastq2 Conversion Software (v. 2.20, Illumina), and primary data analysis was performed with the Cell Ranger Gene Expression pipeline (version 3.1.0, 10X Genomics).

### Individual sample processing

Data processing was implemented similarly for the three conditions (*i.e.*, fed, 12h-fasting, and 24h-fasting) using R (version 4.1.2). Quality control and preprocessing were performed using SingleCellExperiment (version 1.14.1) to construct the object with barcodes, features, and unique molecular identifiers (UMI) matrix. To retrieve a maximum number of high-quality cells, emptyDrops (dropletUtils version 1.12.3) was applied with default parameters to remove empty barcodes. Additionally, the percentage of reads mapping to the mitochondrial genome is usually used as a proxy to remove dead/dying cells. However, a recent publication^59^ demonstrated that the fraction of mitochondrial reads depends on the tissue of origin, cell type, and experimental conditions: the usual strict cut-off of 5% could eliminate a large proportion of “valid” cells. As these variations may occur in our experimental design (*i.e.*, fed *vs.* fasted conditions), isOutiler used with default parameters (scuttle version 1.2.1) was preferred to arbitrary cut-offs. Briefly, a cell is removed if the value of the mitochondrial expression is 3 MADs (median absolute deviation, the default) away from the median of expression in the population. Finally, to detect and remove putative doublet cells, scDblFinder (version 1.6.0) was employed, and cells with a doublet score >5 (arbitrary) were removed from the analysis. Before data normalization with Seurat (version 4.0.6), a last quality control step was added to remove all cells with less than 200 genes detected.

### Clustering analysis

Primary processing was done with Seurat using quality-controlled cells as input. Data normalization was performed with SCTransform() followed by Principal Component decomposition (RunPCA(), default parameter) and non-linear dimension reduction (RunUMAP(), with n_dim dimensions, where n_dim is selected as follows: the number of dimensions corresponding to the 3^rd^ quantile of the explained variance distribution rather than being set arbitrarily based on the visual inspection of ElbowPlot()). Clustering was performed from the UMAP representation by calculating a Share Nearest Neighbor graph (FindNeighbors()0) and applying a modularity-based cluster detection (FindClusters() with default parameters.

Cell-type assignment per cluster was done by *i.* extracting markers (FindMarkers()) and *ii.* comparing them with current knowledge and literature revision. To identify the biological processes characteristics for each cluster, enrichment analysis was performed with enrichR (version 3.1) using the most recent databases for “Biological Process (2021)”, “Cellular Component (2021)”, “KEGG (2019)”, and “GWAS Catalog 2019“.

### Pseudospatial analysis

To identify the pseudospatial gradual patterns along the 3V, normalization, dimensional reduction, and clustering were performed in the feeding condition as described above. Note that in our dataset, ependyma cell populations are clustered in an anatomically-like distribution. The resulting Seurat object was converted to CellDataSet using the *as.cell_data_set* function of *Seurat-wrapper.* Following conversion, we used Monocle3’s functions *learn_graph()* and *order_cells()* with default parameters to order cells based on their pseudospatial trajectory from ependyma to β2 tanycytes. To identify genes highly expressed to each ependyma cell subgroup (*e.g.*, Epen, α1, β1) as specific markers *versus* genes spanning several populations (*e.g.*, epen1+α1, α1+β1) as overlapping markers, we calculated the correlation between each gene expression vector and binary vectors representing simple or combined populations (1 if the cell belongs to the population, 0 otherwise). Correlations were performed using the *rcorr* function (Hmisc v. 5.01). Only significant (p.adjusted<0.01) and positive correlations were considered. Then, for each gene, a score in the form of the product of the correlation and its weighted average expression was calculated to normalize for gene expression levels. A gene was classified as specific to a given ependyma population or belonging to a combined population according to the highest score among all possible classifications. Pseudospatial trajectories were visualized using a cubic spline-based regression on pseudospatial gradients and expression levels.

### Integrated analysis

All single cells datasets were integrated together^60^ to compare clusters between the different metabolic conditions (Fed, 12h-Fasting, and 24h-Fasting) in a common background. Quality control and data normalization were done independently for each sample with the same parameters as above. For the integration, datasets are processed with SelectIntegrationFeatures(), PrepSCTIntegration(), FindIntegrationAnchors(), and IntegrateData() with default parameters. These functions find variable features common between all datasets, identify anchors (pair of cells from each dataset determined as a mutual nearest neighbor (MNN)), and integrate datasets based on filtered anchors^60^. The further processing is the same as the individual sample processing.

### Differential gene expression analysis

The integrated object with the three metabolic conditions was used to perform differential gene expression analysis. Default Seurat workflow was questioned by a recent publication^61^ showing many false positive differentially expressed genes influencing downstream analyses. Thus, we designed an alternate workflow based on pseudobulks, taking advantage of the proven reliability of the DESeq2 package (version 1.38.3). Briefly, each dataset (*i.e.*, the three conditions) was split into four pseudo-replicates with an equivalent number of cells per cluster, and DESeq2 with default parameter was applied.

### Differential pseudospatial analysis

To determine the transcriptional gradient between ependymal cell populations and identify their differences across conditions, we perform a differential pseudospatial analysis. Using the integrated Seurat object, with normalized and integrated counts, we calculated a gene expression correlation matrix for the 12h-fasting and 24h-fasting conditions as described below. Tradeseq was used to calculate pseudospatial patterns using default parameters. The number of patterns was subsetted at those with more than 50 genes.

### Transcription factors activity estimation

We used pySCENIC v.0.12.1 to process the transcriptional profile of all known transcription factors in the three metabolic conditions. Transcription factors activity has been estimated using the AUCell module from the regulons obtained from the standard GNRBoost processing of the co-expression matrix. The heatmap has been generated by clustering on the columns (Genes) only (parameter Rowv = NA) to keep the pseudospatial ordering on the rows.

### Inference of tanycyte-neuron communications

To infer mechanisms of cell-cell communication between tanycytes and neighboring subpopulations in the scRNAseq, we generated a method based on the expression product of ligand-receptor interactions. To infer cell-cell interactions, we used the ligand-receptor pairs database from CellChat^62^. Using the integrated Seurat object, we subset each condition (*i.e.*, fed, 12h-fasting, and 24h-fasting). For each ligand-receptor pair, a function calculates Communication Scores based on the average of the product of expression from each cell of one cluster (expressing the ligand) to each of the other clusters (expressing the receptor). An ANOVA was calculated on the matrix of communication scores for a given ligand-receptor pair on the interaction between two cell types to determine the differences between conditions.

### Histology

TAT-Cre-injected tdTomato mice were anesthetized with isoflurane and perfused transcardially with 0.9% saline, followed by an ice-cold solution of 4% paraformaldehyde in 0.1 M phosphate buffer (pH 7.4). Brains were quickly removed, postfixed in the same fixative for two hours at 4°C, and immersed in 20% sucrose in 0.1M phosphate-buffered saline (PBS) at 4°C overnight. Brains were finally embedded in an ice-cold OCT medium (optimal cutting temperature embedding medium, Tissue Tek, Sakura) and frozen on liquid nitrogen-cooled isopentane. Brains were cut using a cryostat into 20-μm-thick coronal sections and processed for staining as described previously^15^. Briefly, slide-mounted sections were washed, counterstained with DAPI (1:10,000, Molecular Probes, Invitrogen), and coverslipped using Mowiol (Calbiochem, La Jolla, CA).

### Data availability

The accession number for the single-cell transcriptome reported in this paper is Gene Expression Omnibus (GEO; www.ncbi.nlm.nih.gov/geo): XXX. R codes are available through GitHub (ZZZ).

