## Supplemental figures for "Pseudospatial transcriptional gradient analysis of hypothalamic ependymal cells: towards a new tanycyte classification"

### Supplemental information

Brunner M.[1,†], Lopez-Rodriguez D.[2,†], Messina A.[1], Thorens B.[3], Santoni F[1,*], Langlet F.[2,*]

[1], Service of Endocrinology, Diabetology, and Metabolism, Lausanne University Hospital, 1011 Lausanne, Switzerland.
[2], Department of Biomedical Sciences, Faculty of Biology and Medicine, University of Lausanne, Lausanne, Switzerland
[3], Center for Integrative Genomics, Faculty of Biology and Medicine, University of Lausanne, Lausanne, Switzerland

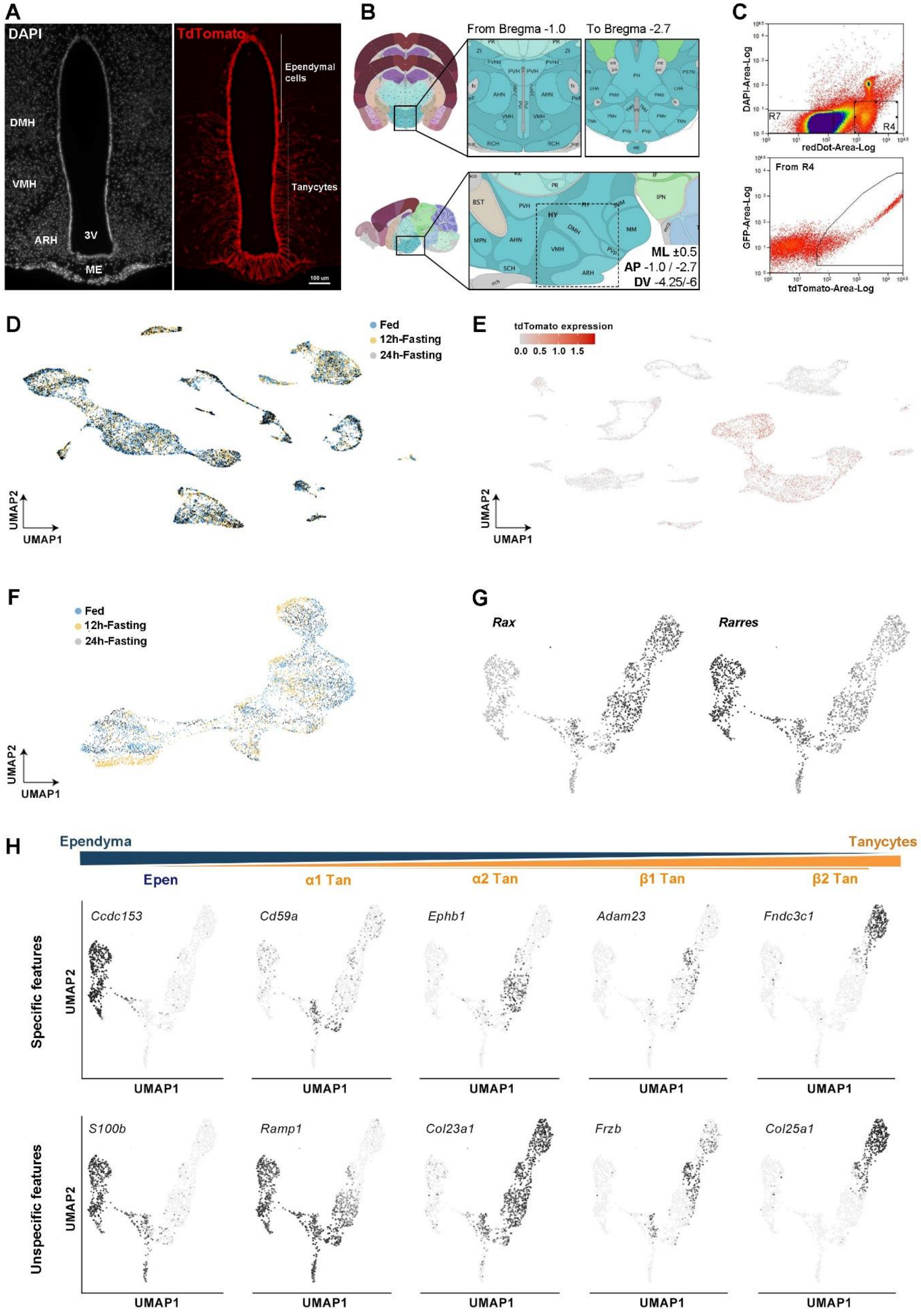
A
DAPI
TdTomato
Ependymal cells
Tanycytes
DMH
VMH
ARH
3V
ME
100 um
B
From Bregma -1.0
To Bregma -2.7
ML ±0.5
AP -1.0 / -2.7
DV -4.25/-6
C
DAPI-Area-Log
redDot-Area-Log
R7
R4
From R4
GFP-Area-Log
tdTomato-Area-Log
D
Fed
12h-Fasting
24h-Fasting
UMAP2
UMAP1
E
tdTomato expression
0.0 0.5 1.0 1.5
UMAP2
UMAP1
F
Fed
12h-Fasting
24h-Fasting
UMAP2
UMAP1
G
Rax
Rarres
H
Ependyma
Tanycytes
Epen
α1 Tan
α2 Tan
β1 Tan
β2 Tan
Specific features
Ccdc153
Cd59a
Ephb1
Adam23
Fndc3c1
Unspecific features
S100b
Ramp1
Col23a1
Frzb
Col25a1
UMAP2
UMAP1

**Supplementary Figure 1. (A)** DAPI counterstaining (left) and tdTomato expression (right) along the 3V after TAT-cre infusion. Tanycytes and ependymal cells were selectively targeted. **(B)** Schematic representation of the microdissected region. **(C)** Gating strategy for fluorescence-activated cell sorting (FACS). Cells were sorted according to their negativity for DAPI (viability dye), their positivity for RedDot 1 (viability dye), and their level of tdTomato fluorescence. **(D)** UMAP representation of the three integrated conditions (*i.e.*, fed, 12h-fasting, and 24h-fasting), colored by metabolic conditions. **(E)** tdTomato expression among sorted cells. **(F)** UMAP representation of the three integrated conditions (*i.e.*, fed, 12h-fasting, and 24h-fasting) for the ependymal clusters, colored by metabolic conditions. **(G)** *Rax* and *Rarres* UMAP gene expression in the ependymal cell populations. **(H)** Gene expression of specific *versus* unspecific features for each ependymal cell subgroup.

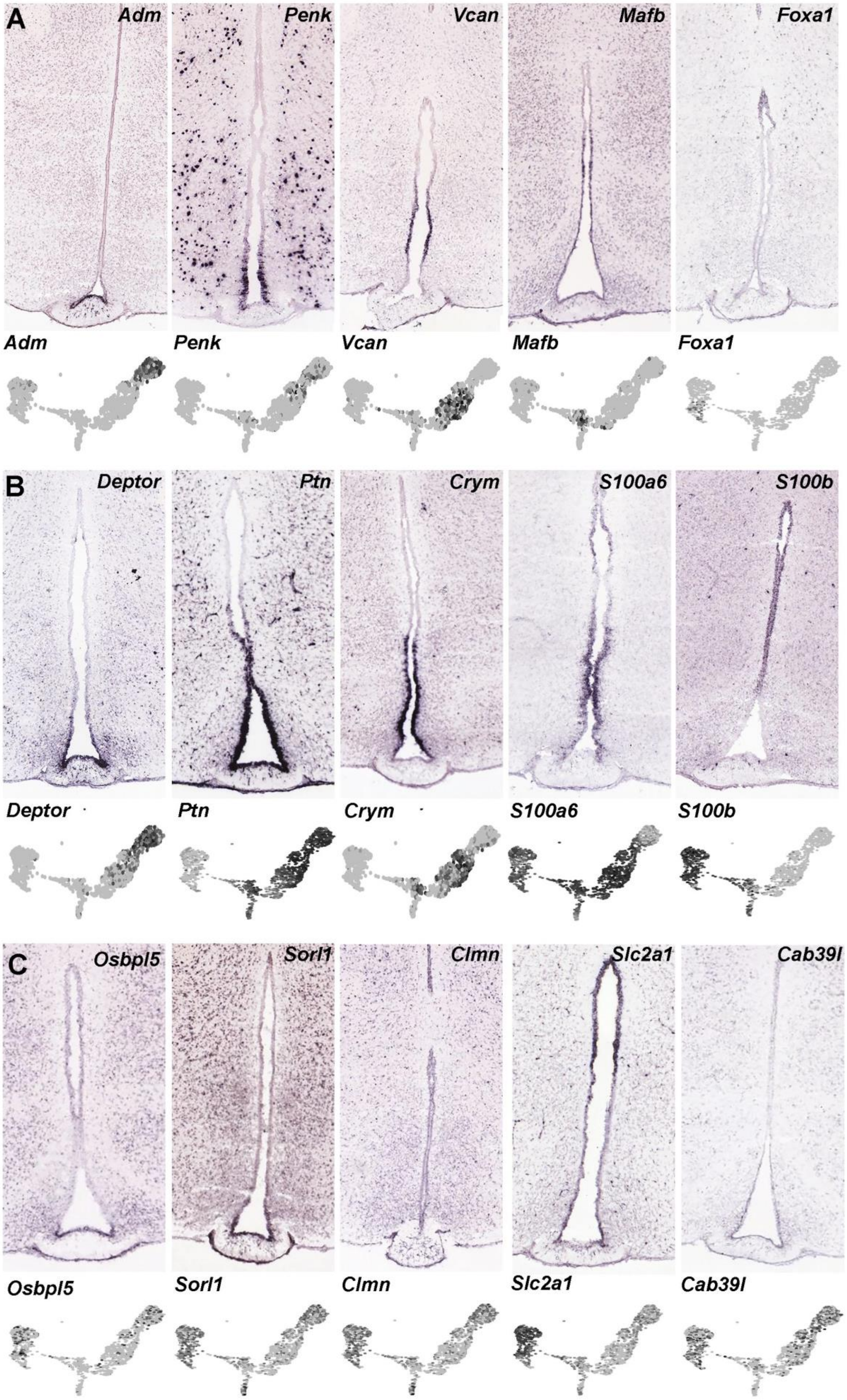
A
Adm
Penk
Vcan
Mafb
Foxa1
Adm
Penk
Vcan
Mafb
Foxa1
B
Deptor
Ptn
Crym
S100a6
S100b
Deptor
Ptn
Crym
S100a6
S100b
C
Osbpl5
Sorl1
Clmn
Slc2a1
Cab39l
Osbpl5
Sorl1
Clmn
Slc2a1
Cab39l

**Supplementary Figure 2. (A)** *In situ* hybridization on coronal brain sections (Allen Mouse Brain Atlas) and feature plot derived from UMAP showing the expression and distribution of different features. Features were selected for their specificity for each cluster. **(B)** *In situ* hybridization on coronal brain sections (Allen Mouse Brain Atlas) and feature plot derived from UMAP showing the expression and distribution of different features. Features were selected from overlapping markers spanning different clusters. **(C)** *In situ* hybridization on coronal brain sections (Allen Mouse Brain Atlas) and feature plot derived from UMAP showing the expression and distribution of different features. Features were selected for their U-shape pattern along the 3V.

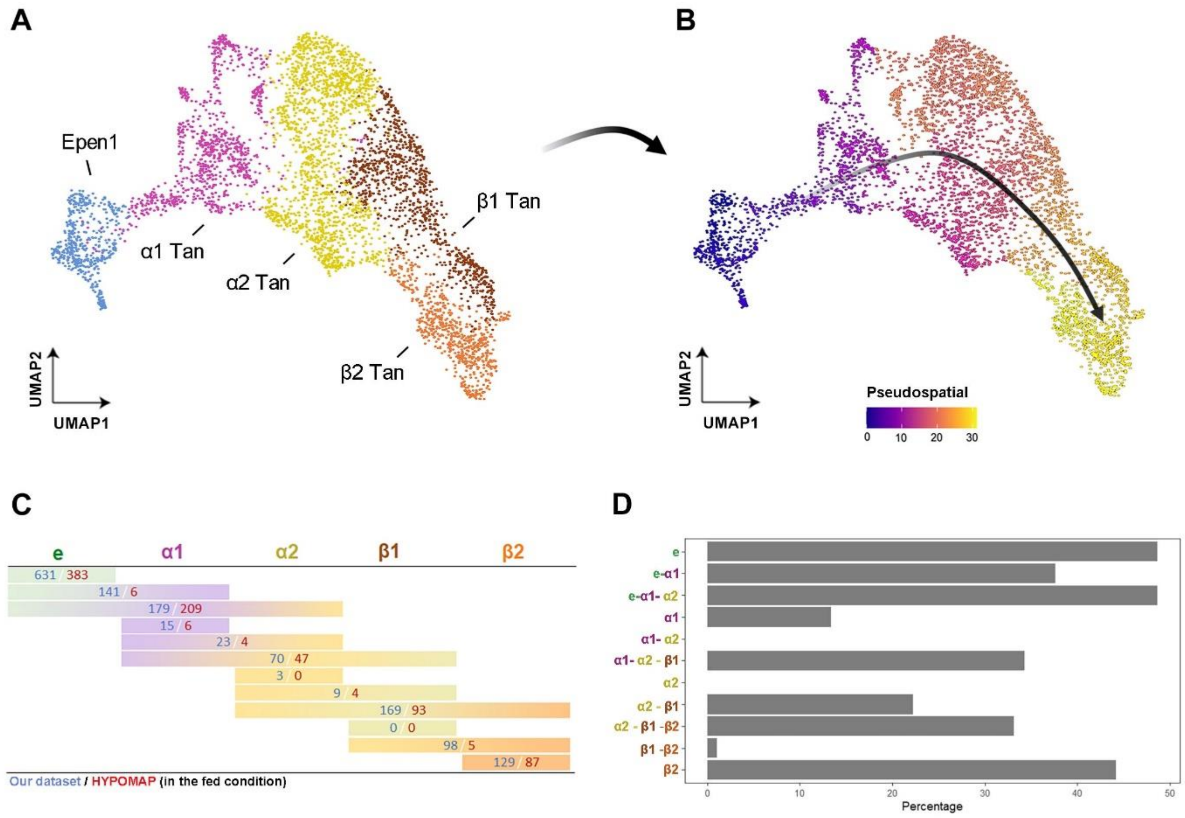


**Supplementary Figure 3. Pseudospatial analysis done on the Hypomap dataset. (A-B)** UMAP representation of hypomap cells in fed conditions (A) converting into a pseudospatial trajectory from typical ependymal cells to β2-tanycytes (B). **(C)** Number of features significantly correlating with one (specific markers) *versus* multiple ependymal populations (overlapping markers) found in our dataset (blue, fed) and hypomap dataset (red, hypomap). **(D)** Percentage of shared features between our dataset and hypomap dataset for each subgroup.

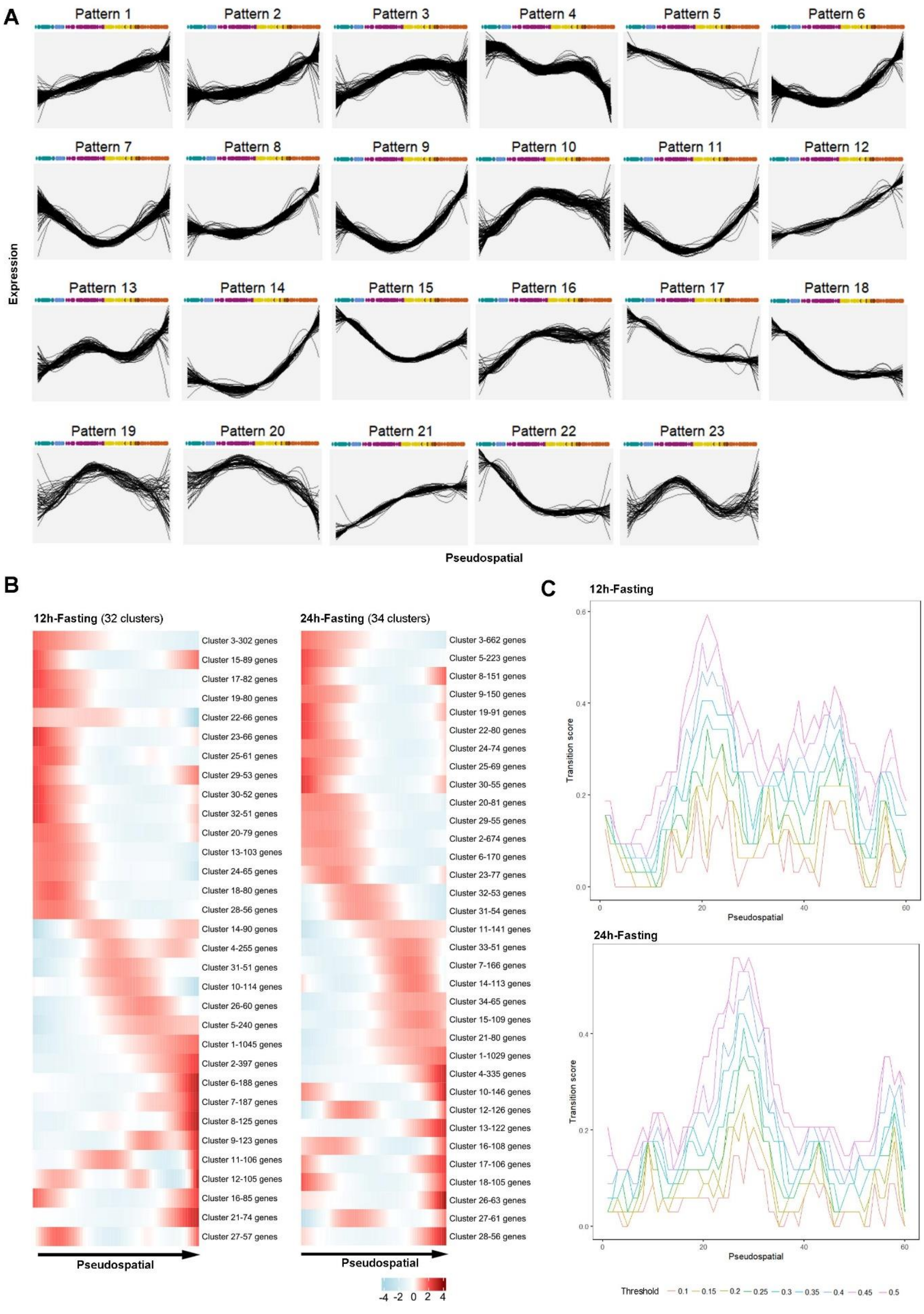
A
Pattern 1
Pattern 2
Pattern 3
Pattern 4
Pattern 5
Pattern 6
Pattern 7
Pattern 8
Pattern 9
Pattern 10
Pattern 11
Pattern 12
Pattern 13
Pattern 14
Pattern 15
Pattern 16
Pattern 17
Pattern 18
Pattern 19
Pattern 20
Pattern 21
Pattern 22
Pattern 23
Expression
Pseudospatial
B
12h-Fasting (32 clusters)
Cluster 3-302 genes
Cluster 15-89 genes
Cluster 17-82 genes
Cluster 19-80 genes
Cluster 22-66 genes
Cluster 23-66 genes
Cluster 25-61 genes
Cluster 29-53 genes
Cluster 30-52 genes
Cluster 32-51 genes
Cluster 20-79 genes
Cluster 13-103 genes
Cluster 24-65 genes
Cluster 18-80 genes
Cluster 28-56 genes
Cluster 14-90 genes
Cluster 4-255 genes
Cluster 31-51 genes
Cluster 10-114 genes
Cluster 26-60 genes
Cluster 5-240 genes
Cluster 1-1045 genes
Cluster 2-397 genes
Cluster 6-188 genes
Cluster 7-187 genes
Cluster 8-125 genes
Cluster 9-123 genes
Cluster 11-106 genes
Cluster 12-105 genes
Cluster 16-85 genes
Cluster 21-74 genes
Cluster 27-57 genes
Pseudospatial
24h-Fasting (34 clusters)
Cluster 3-662 genes
Cluster 5-223 genes
Cluster 8-151 genes
Cluster 9-150 genes
Cluster 19-91 genes
Cluster 22-80 genes
Cluster 24-74 genes
Cluster 25-69 genes
Cluster 30-55 genes
Cluster 20-81 genes
Cluster 29-55 genes
Cluster 2-674 genes
Cluster 6-170 genes
Cluster 23-77 genes
Cluster 32-53 genes
Cluster 31-54 genes
Cluster 11-141 genes
Cluster 33-51 genes
Cluster 7-166 genes
Cluster 14-113 genes
Cluster 34-65 genes
Cluster 15-109 genes
Cluster 21-80 genes
Cluster 1-1029 genes
Cluster 4-335 genes
Cluster 10-146 genes
Cluster 12-126 genes
Cluster 13-122 genes
Cluster 16-108 genes
Cluster 17-106 genes
Cluster 18-105 genes
Cluster 26-63 genes
Cluster 27-61 genes
Cluster 28-56 genes
Pseudospatial
-4 -2 0 2 4
C
12h-Fasting
Transition score
Pseudospatial
24h-Fasting
Transition score
Pseudospatial
Threshold 0.1 0.15 0.2 0.25 0.3 0.35 0.4 0.45 0.5

**Supplementary Figure 4. (A)** Twenty-three patterns of gene expression observed along the pseudospatial trajectory obtained using TradeSeq. **(B)** Heatmap showing different shared gene expression patterns along the pseudospatial trajectory in 12h-fasting and 24h-fasting. **(C)** Graph representing the main transition region along the pseudospatial trajectory. The transition score was calculated as an on-off switch in gene expression along the trajectory. Values represent on-off expression gating thresholds.

**Supplementary Table S1. A.** Experiment summary (related to Fig.1A). **B.** Cell count on the general clustering workflow (related to Fig.1B). **C.** Number of tdTomato-expressing cells (related to SuppFig.1). **D.** Cell count on the ependyma clustering workflow (related to Fig.1D).

**Supplementary Table S2. A.** List of features for each cluster on the general clustering workflow (related to Fig.1C). **B.** List of features for each cluster on the ependyma clustering workflow in the fed condition (related to Fig.1E). **C-G.** GO_MolecularFunction (C), GO_BiologicalProcess (D), GO_CellularComponent (E), KEGG (F), and GWAS enrichment (G) for each cluster on the ependyma clustering workflow in the fed condition (related to Fig.1F).

**Supplementary Table S3. A.** List of specific *versus* overlapping features for each ependymal population in the fed, 12h-fasting, and 24h-fasting conditions (related to Fig.2C and 5B). **B-D.** GO_MolecularFunction (MF), GO_BiologicalProcess (BP), GO_CellularComponent (CC), KEGG, and GWAS enrichment for each group along the ependyma in the fed condition (B, related to Fig. 2C), 12h-fasting (C), and 24h-fasting (D) conditions. **E.** Percentage of co-expressing cells for each tanycyte population in the fed condition (related to Fig.2D).

**Supplementary Table S4. A.** List of features for each cluster on the ependyma clustering workflow for the fed, 12h-fasting, and 24h-fasting conditions separately (related to Fig. 3A). **B.** Number and percentage of shared markers between subgroups and metabolic conditions, used to build the differentiate tree (related to Fig. 3C). **C.** List of features for each cluster on the ependyma clustering workflow for the integrated fed, 12h-fasting, and 24h-fasting conditions (related to Fig. 3C). **D-F.** List of differentially expressed genes for each cluster in Fed *vs*. 12h-fasting (D), Fed *vs.*

24h-fasting (E), and 12h-fasting *vs.* 24h-fasting (F) on the ependyma clustering workflow (related to Fig. 3E).

**Supplementary Table S5. A.** List of features for each temporal trajectory from fed to 24h-fasting (related to Fig. 4B). **B-F.** GO_MolecularFunction (B), GO_BiologicalProcess (C), GO_CellularComponent (D), KEGG (E), and GWAS enrichment (F) for each temporal trajectory on the ependyma clustering workflow (related to Fig. 4C).

**Supplementary Table S6. A.** List of features in the twenty-three ventrodorsal pattern along the third ventricle in the fed condition. **B.** GO_MolecularFunction (MF), GO_BiologicalProcess (BP), GO_CellularComponent (CC), KEGG, and GWAS enrichment for each pattern along the ependyma (related to Fig. 5B).

**Supplementary Table S7. A-D.** List of differentially regulated ligand-receptor couples in fed *vs.* 12h-fasting and fed *vs.* 24h-fasting conditions between β1 tanycytes and vessels (A), β1 tanycytes and neurons (B), β2 tanycytes and vessels (C), and β2 tanycytes and neurons (D). **E.** Number and percentage of cells co-expressing specific ligands for BBB contact *versus* fenestrated vessels contact. The high percentage means that tanycytes express specific ligands to contact BBB and fenestrated vessels.
